# Likelihood Approximation Networks (LANs) for Fast Inference of Simulation Models in Cognitive Neuroscience

**DOI:** 10.1101/2020.11.20.392274

**Authors:** Alexander Fengler, Lakshmi N. Govindarajan, Tony Chen, Michael J. Frank

## Abstract

In cognitive neuroscience, computational modeling can formally adjudicate between theories and affords quantitative fits to behavioral/brain data. Pragmatically, however, the space of plausible generative models considered is dramatically limited by the set of models with known likelihood functions. For many models, the lack of a closed-form likelihood typically impedes Bayesian inference methods. As a result, standard models are evaluated for convenience, even when other models might be superior. Likelihood-free methods exist but are limited by their computational cost or their restriction to particular inference scenarios. Here, we propose neural networks that learn approximate likelihoods for arbitrary generative models, allowing fast posterior sampling with only a one-off cost for model simulations that is amortized for future inference. We show that these methods can accurately recover posterior parameter distributions for a variety of neurocognitive process models. We provide code allowing users to deploy these methods for arbitrary hierarchical model instantiations without further training.

## 1 Introduction

Computational modeling has gained traction in cognitive neuroscience in part because it can guide principled interpretations of functional demands of cognitive systems while maintaining a level of tractability in the production of quantitative fits of brain-behavior relationships. For example, models of reinforcement learning are frequently used to estimate the neural correlates of the exploration/exploitation tradeoff, of asymmetric learning from positive versus negative outcomes, or of model-based vs model-free control (***Schönberg et al., 2007***; ***Niv et al., 2012***; ***Frank et al., 2007***; ***Zajkowski et al., 2017***; ***Badre et al., 2012***; ***Daw et al., 2011b***). Similarly, models of dynamic decision-making processes are commonly used to disentangle the strength of the evidence for a given choice from the amount of that evidence needed to commit to any choice, and how such parameters are impacted by reward, attention, and neural variability across species (***Rangel et al., 2008***; ***Forstmann et al., 2010***; ***Krajbich and Rangel, 2011***; ***Frank et al., 2015***; ***Yartsev et al., 2018***; ***Doi et al., 2020***). Parameter estimates might also be used as a theoretically-driven method to reduce the dimensionality of brain/behavioral data that can be used for prediction of e.g. clinical status in computational psychiatry (***Huys et al., 2016***).

Interpreting such parameter estimates requires robust methods that can estimate their generative values, ideally including their uncertainty. For this purpose, Bayesian statistical methods have gained traction. The basic conceptual idea in Bayesian statistics is to treat parameters *θ* and data **x** as stemming from a joint probability model *p*(*θ,* **x**). Statistical inference proceeds by using Bayes’ rule,

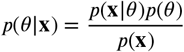

to get at *p*(*θ*|**x**), the posterior distribution over parameters given data. *p*(*θ*) is known as the prior distribution over the parameters *θ*, and *p*(**x**) is known as the evidence or just probability of the data (a quantity of importance for model comparison). Bayesian parameter estimation is a natural way to characterize uncertainty over parameter values. In turn, it provides a way to identify and probe parameter trade-offs. While we often don’t have access to *p*(*θ*|**x**) directly, we can draw samples from it instead, for example via Markov Chain Monte Carlo (***Robert and Casella, 2013***; ***Diaconis, 2009***; ***Robert and Casella, 2011***).

Bayesian estimation of the full posterior distributions over model parameters contrast with maximum likelihood estimation methods often used to provide single best parameter values, without considering their uncertainty or whether other parameter estimates might give similar fits. Bayesian methods naturally extend to settings that assume an implicit hierarchy in the generative model in which parameter estimates at the individual level are informed by the distribution across a group, or even to assess within an individual how trial-by-trial variation in (for example) neural activity can impact parameter estimates (commonly known simply as hierarchical inference). Several toolboxes exist for estimating the parameters of popular models like the drift diffusion model of decision making and are widely used by the community for this purpose (***Wiecki et al., 2013***; ***Heathcote et al., 2019***; ***Turner et al., 2015a***; ***Ahn et al., 2017***). Various studies have used these methods to characterize how variability in neural activity, and manipulations thereof, alter learning and decision parameters that can quantitatively explain variability in choice and response time distributions (***Cavanagh et al., 2011***; ***Frank et al., 2015***; ***Herz et al., 2016***; ***Pedersen and Frank, 2020***).

Traditionally, however, posterior sampling or maximum likelihood estimation for such models required analytical likelihood functions: a closed-form mathematical expression providing the likelihood of observing specific data (reaction times and choices) for a given model parameterization. This requirement limits the application of any likelihood-based method to a relatively small subset of cognitive models chosen for so-defined convenience instead of theoretical interest. Consequently, model comparison and estimation exercises are constrained, as many important but likelihood-free models were effectively *untestable* or required weeks to process a single model formulation. Testing any slight adjustment to the generative model (e.g., different hierarchical grouping or splitting conditions) requires a repeated time investment of the same order. For illustration, we focus on the class of sequential sampling models applied to decision-making scenarios, with the most common variants of the drift diffusion model (DDM). The approach is, however, applicable to any arbitrary domain.

In the standard DDM, a two-alternative choice decision is modeled as a noisy accumulation of evidence toward one of two decision boundaries (***Ratcliff and McKoon, 2008***). This model is widely used as it can flexibly capture variations in choice, error rates, and response time distributions across a range of cognitive domains and its parameters have both psychological and neural implications. While the likelihood function is available for the standard DDM and some variants including inter-trial variability of its drift parameter, even seemingly small changes to the model form, such as dynamically varying decision bounds (***Cisek et al., 2009***; ***Hawkins et al., 2015***) or multiple choice alternatives (***Krajbich and Rangel, 2011***), are prohibitive for likelihood-based estimation, and instead require expensive monte carlo simulations, often without providing estimates of uncertainty across parameters.

In the last decade and a half, Approximate Bayesian Computation (ABC) algorithms have grown to prominence (***Sisson et al., 2018***). These algorithms enable one to sample from posterior distributions over model parameters, where models are defined only as simulators, which can be used to construct empirical likelihood distributions (***Sisson et al., 2018***). ABC approaches have enjoyed successful application across life and physical sciences (e.g., ***Akeret et al. (2015)***), and notably, in cognitive science (***Turner and Van Zandt, 2018***), enabling researchers to estimate theoretically interesting models that were heretofore intractable. However, while there have been many advances without sacrificing information loss in the posterior distributions (***Turner and Sederberg, 2014***; ***Holmes, 2015***), such ABC methods typically require many simulations to generate synthetic or empirical likelihood distributions, and hence can be computationally expensive (in some cases prohibitive – it can take weeks to estimate parameters for a single model). This issue is further exacerbated when embedded within a sequential Markov Chain Monte Carlo (MCMC) sampling scheme, which is needed for unbiased estimates of posterior distributions. For example, one typically needs to simulate between 10,000 - 100,000 times (the exact number varies depending on the model) for each proposed combination of parameters (i.e., for each sample along a Markov chain, which may itself contain tens of thousands of samples), after which they are discarded. This situation is illustrated in Figure 1, where the red arrows point at the computations involved in the approach suggested by ***Turner et al. (2015b)***.

**Figure 1.**
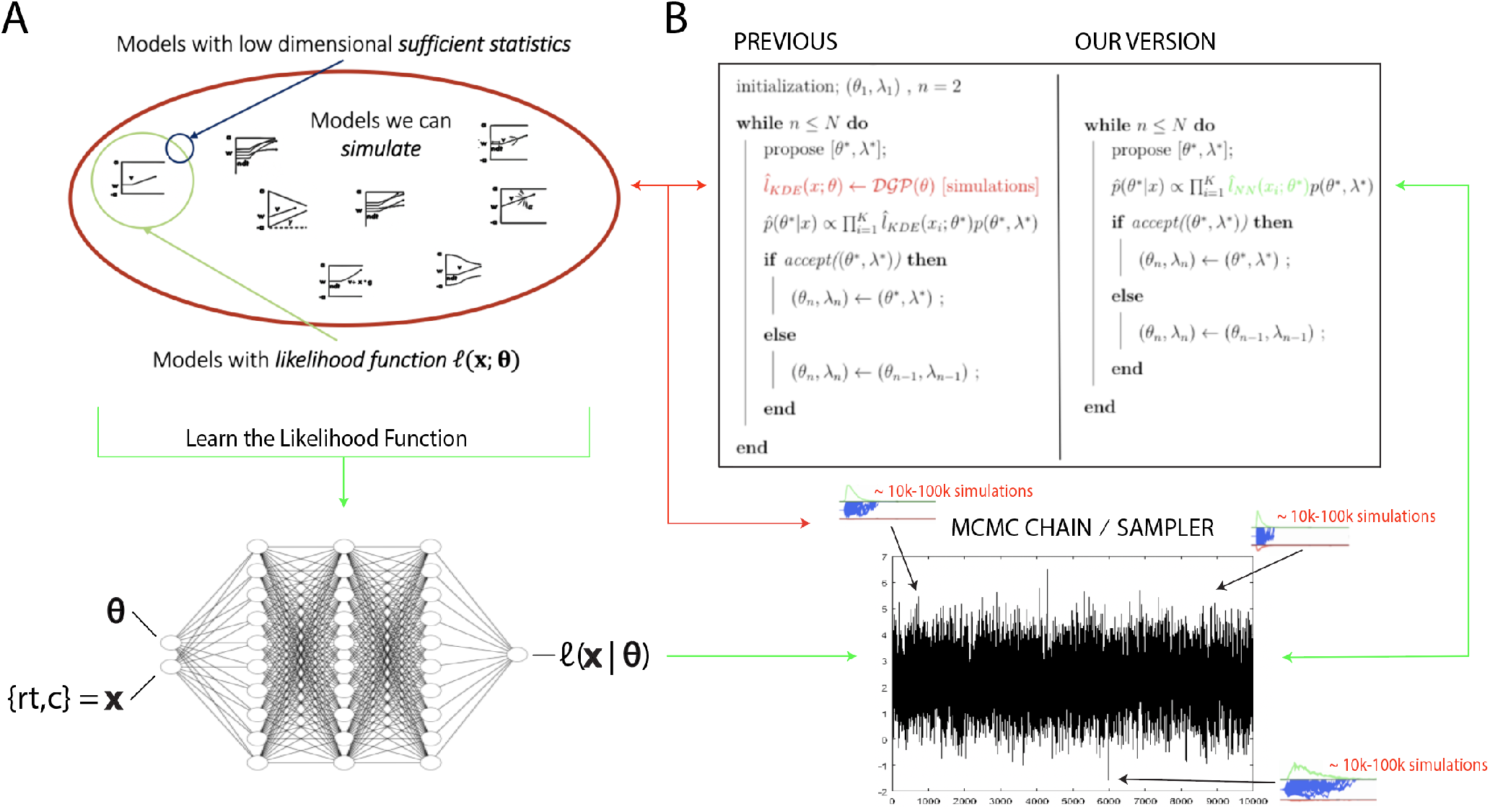
**A** The space of theoretically interesting models in the cognitive neurosciences (red) is much larger than the space of mechanistic models with analytic likelihood functions green. Traditional ABC methods require models that have low dimensional sufficient statistics (blue). **B** Illustrates how LANs can be used in lieu of online simulations for efficient posterior sampling. The left panel shows the predominant PDA method used for ABC in the cognitive sciences (***Turner et al., 2015b***). For each step along a Markov chain, 10K-100K simulations are required to obtain a single likelihood estimate. The right panel shows how we can avoid the simulation steps during inference using amortized likelihood networks that have been pretrained using PDAs.

To address this type of issue, the statistics and machine learning communities have increasingly focused on strategies for the amortization of simulation-based computations (***Gutmann et al., 2018***; ***Papamakarios and Murray, 2016***; ***Papamakarios et al., 2019a***; ***Lueckmann et al., 2019***; ***Radev et al., 2020a, b***; ***Goncalves et al., 2020***). The aim is generally to use model simulations up-front and learn a reusable approximation of the function of interest (targets can be the likelihood, the evidence, or the posterior directly).

In this article, we develop a general ABC method (and toolbox) that allows users to perform inference on a significant number of neurocognitive models without repeatedly incurring substantial simulation costs. To motivate our particular approach and situate it in the context of other methods, we outline the following key desiderata:

1. First, the method needs to be easily and rapidly deployable to end-users for Bayesian inference on various models hitherto deemed computationally intractable. This desideratum naturally leads us to an amortization approach, where end-users can benefit from costs incurred upfront.
2. Second, our method should be sufficiently flexible to support arbitrary inference scenarios, including hierarchical inference, estimation of latent covariate (e.g., neural) processes on the model parameters, arbitrary specification of parameters across experimental conditions, and without limitations on data-set sizes. This desideratum leads us to amortize the likelihood functions, which (unlike other amortization strategies) can be immediate applied to arbitrary inference scenarios without further cost.
3. Third, we desire approximations that do not apriori sacrifice covariance structure in the parameter posteriors, a limitation often induced for tractability in variational approaches to approximate inference (***Blei et al., 2017***).
4. Fourth, end-users should have access to a convenient interface that integrates the new methods seamlessly into pre-existing workflows. The aim is to allow users to get access to a growing database of amortized models through this toolbox and enable increasingly complex models to be fit to experimental data routinely, with minimal adjustments to the user’s working code. For this purpose, we will provide an extension to the widely used HDDM toolbox (***Wiecki et al., 2013***; ***Pedersen and Frank, 2020***) for parameter estimation of DDM and RL models.
5. Fifth, our approach should exploit modern computing architectures, specifically parallel computation. This leads us to focus on the encapsulation of likelihoods into neural network architectures, which will allow batch-processing for posterior inference.

Guided by these desiderata, we developed two amortization-based algorithms using Neural Networks (NNs) in combination with KDE. Rather than simulate during inference, we instead train NNs as parametric function approximators to learn the likelihood function from an initial set of *apriori* simulations across a wide range of parameters. By learning likelihood functions directly, we avoid posterior distortions that result from inappropriately chosen (user defined) summary statistics and corresponding distance measures as applied in traditional ABC methods (***Sisson et al., 2018***). Once trained, likelihood evaluation only requires a forward pass through the NN (as if it were an analytic likelihood) instead of necessitating simulations. Moreover, any algorithm can be used to facilitate posterior sampling, maximum likelihood estimation (MLE), or maximum a posteriori estimation (MAP).

For generality, and because they each have their advantages, we use two classes of architectures, Multi-layered-perceptrons (MLPs) and Convolutional Neural Networks (CNNs), and two different posterior sampling methods (MCMC and Importance sampling). We show proofs of concepts using posterior sampling and parameter recovery studies for a range of cognitive process models of interest. The trained neural networks provide the community with a (continually expandable) bank of encapsulated likelihood functions that facilitate consideration of a larger (previously computationally inaccessible) set of cognitive models, with orders of magnitude speedup relative to simulation-based methods. This speedup is possible because costly simulations only have to be run once per model upfront and henceforth be avoided during inference: previously executed computations are amortized and then shared with the community.

Moreover, we develop standardized amortization-pipelines that allow the user to apply this method to arbitrary models, requiring them to provide only a functioning simulator of their model of choice.

In Section 2 we situate our approach in the greater context of approximate Bayesian computation (ABC), with a brief review of online and amortization algorithms. Section 3 describes the amortization-pipelines we propose as well as suggested use-cases. Section 4 discusses the cognitive process models that we utilized as relevant test-beds for our methods. Section 5 shows proof-of-concept parameter recovery studies for the two proposed algorithms, demonstrating that the method accurately recovers both the posterior mean and variance (uncertainty) of generative model parameters, and that it does so at a run-time speed of orders of magnitude faster than traditional ABC approaches without further training. In Section 6 we further demonstrate an application to hierarchical inference, in which our trained networks can be imported into widely used toolboxes for arbitrary inference scenarios. In Section 7 and Section 8, we further situate our work in the context of other ABC amortization strategies and discuss limitations and future work.

## 2 Approximate Bayesian Computation

ABC methods apply when one has access to a parametric stochastic simulator (also referred to as generative model), but, unlike the usual setting for statistical inference, no access to an explicit mathematical formula for the likelihood of observations given the simulator’s parameterization.

While likelihood functions for a stochastic stimulator can in principle be mathematically derived, this can be exceedingly challenging even for some historically famous models such as the Ornstein-Uhlenbeck process (***Lipton and Kaushansky, 2018***)), and maybe intractable in many others. Consequently, the statistical community has increasingly developed ABC tools that enable posterior inference of such “likelihood-free” stochastic models, while completely bypassing any likelihood derivations (***Cranmer et al., 2020***).

Given a parametric stochastic simulator model 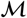, instead of exact inference based on 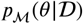, these methods attempt to draw samples from an approximate posterior 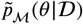. Consider the following general equation:

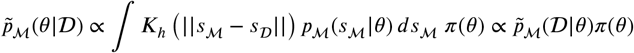

where 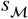 refers to sufficient statistics (roughly, summary statistics of a dataset that retain sufficient information about the parameters of the generative process).^1^ 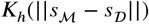, refers to a kernel-based distance measure / cost function, which evaluates a probability density function for a given distance between the observed and expected summary statistics 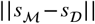. The parameter *h* (commonly known as bandwidth parameter) modulates the cost gradient. Higher values of *h* lead to more graceful decreases in cost (and therefore a worse approximation of the true posterior).

By generating simulations, one can use such summary statistics to obtain approximate likelihood functions, denoted as 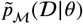, where approximation error can be mitigated by generating large numbers of simulations. The caveat is that the amount of simulation runs needed to achieve a desired degree of accuracy in the posterior can render such techniques computationally infeasible.

With a focus on amortization, our goal is to leverage some of the insights and developments in the ABC community to develop neural network architectures that can learn approximate likelihoods deployable for any inference scenario (and indeed any inference method, including MCMC, variational inference, or even maximum likelihood estimation) without necessitating repeated training. We next describe our particular approach, and we return to a more detailed comparison to existing methods in the Discussion.

## 3 Learning the Likelihood with simple Neural Network Architectures

In this section we outline our approach to amortization of computational costs of large numbers of simulations required by traditional ABC methods. Amortization approaches, incur a one-off (potentially quite large) simulation cost to enable cheap, repeated inference for any dataset. Recent research has lead to substantial developments in this area (***Cranmer et al., 2020***). The most straightforward approach is to simply simulate large amounts of data and compile a database of how model parameterizations are related to observed summary statistics of the data (***Mestdagh et al., 2019***). This database can then be used during parameter estimation in empirical datasets using a combination of nearest neighbor search and local interpolation methods. However, this approach suffers from the curse of dimensionality with respect to storage demands (a problem that is magnified with increasing model parameters). Moreover, its reliance on summary statistics (***Sisson et al., 2018***) does not naturally facilitate flexible re-use across inference scenarios (e.g. hierarchical models, multiple conditions while fixing some parameters across conditions etc.).

To fulfill all desiderata outlined in the introduction, we focus on directly encapsulating the likelihood function over empirical observations (i.e., choices. RTs) of a simulation model so that likelihood evaluation is substantially cheaper than constructing (empirical) likelihoods via model simulation online during inference. This strategy then allows for flexible reuse of such approximate likelihood functions 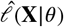, in a large variety of inference scenarios applicable to common experimental design paradigms. Specifically, we encapsulate 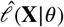 as a feed-forward neural network, which allows for parallel evaluation by design. We refer to these networks as likelihood approximation networks (LANs).

Figure 1, spells out the setting (panel **A**) and usage (panel **B**) of such a method. The architectures for LANs we use in this paper simple, small in size and are made available for download for local usage. While this approach does not allow for instantaneous posterior inference, it does considerably reduce computation time (by up to three orders of magnitude; see “Run-time” section below) when compared to approaches that demand simulations at inference. At the same time we maintain the high degree of flexibility with regards to deployment across arbitrary inference scenarios. As such, LANs can be treated as a highly flexible plugins to existing inference algorithms and remain conceptually simple and lightweight.

Before elaborating our LAN approach, we briefly situate it in the context of more ambitious approaches attempting to amortize posterior inference directly in end-to-end neural networks (***Radev et al., 2020a, b***; ***Papamakarios and Murray, 2016***; ***Papamakarios et al., 2019a***; ***Goncalves et al., 2020***). Here, neural network architectures are trained with a large number of simulated datasets to produce posterior distributions over parameters, and once trained, such networks can be applied to directly estimate parameters from new datasets without need for further simulations. However, the goal to directly estimate posterior parameters from data requires the user to first train a neural network for the very specific inference scenario in which it is applied empirically. Moreover, such approaches are not easily deployable if a user wants to test multiple inference scenarios (e.g., parameters may vary as a function of task condition or brain activity, or in hierarchical frameworks) and consequently, they do not achieve our second desideratum needed for a user-friendly toolbox making inference cheap and simple. We return to discuss the relative merits and limitations of these alternative approaches in the Discussion.

Formally, we use model simulations to learn a function *f*_*w*_ (*x, θ*), where *f*_*w*_ (.) is the output of a neural network with parameter vector **W**(weights, biases). The function *f*_*w*_ (**x***, θ*) is used as an approximation 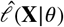 of the likelihood function 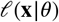. Once learned, we can use such *f* as a plug-in to Bayes Rule,

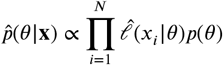

To perform posterior inference we can now use a broad range of existing monte carlo (MC) or markov chain monte carlo (MCMC) algorithms. We note that, fulfilling our second desideratum, an extension to hierarchical inference is as simple as plugging in our neural network into a probabilistic model of the form,

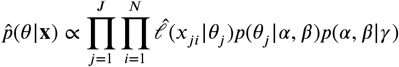

where *α*, *β* are generic group level global parameters, and *γ* serves as a generic fixed hyperparameter vector.

We provide proofs of concepts for two types of LANs (later referred to as MLP, and CNN approaches). The basic difference between these two LANs lies in the problem representation they tackle, while the resulting architectures, respectively multi-layered perceptrons (MLPs) and convolutional neural networks (CNNs), can be treated as naturally falling out of these different problem representations. The first representation, which we call the pointwise approach, considers the functions 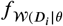, where *θ* is the parameter vector of a given stochastic simulator model, and *D*_*i*_ is a single data point (trial outcome). The pointwise approach is a mapping from the input dimension |*P*|+|*D*|, where *P* refers to the parameters of the simulator model and *D* to the dimensionality of the data generated from this simulator, to the output dimension 1, where the output is simply the likelihood of this single datapoint *D*_*i*_ given the parameterization *P*. As explained in the next section, for this mapping we chose MLPs. The second representation, which we will refer to as the histogram approach, instead aims to learn the discretized global likelihood of an entire dataset for a given parameterization. We represent the output space as an outcome histogram with dimensions *n* × *m* and learn a mapping from input dimension *P* to output dimension *m* × *n*. Representing the problem this way naturally leads to CNNs as the network architecture.

The next two sections will give some detail regarding the networks chosen for the pointwise and histogram LANs. Figure 1 illustrates the general idea of our approach while Figure 2 gives a conceptual overview of the exact training and inference procedure proposed. These details are thoroughly discussed in the last section of the paper.

**Figure 2.**
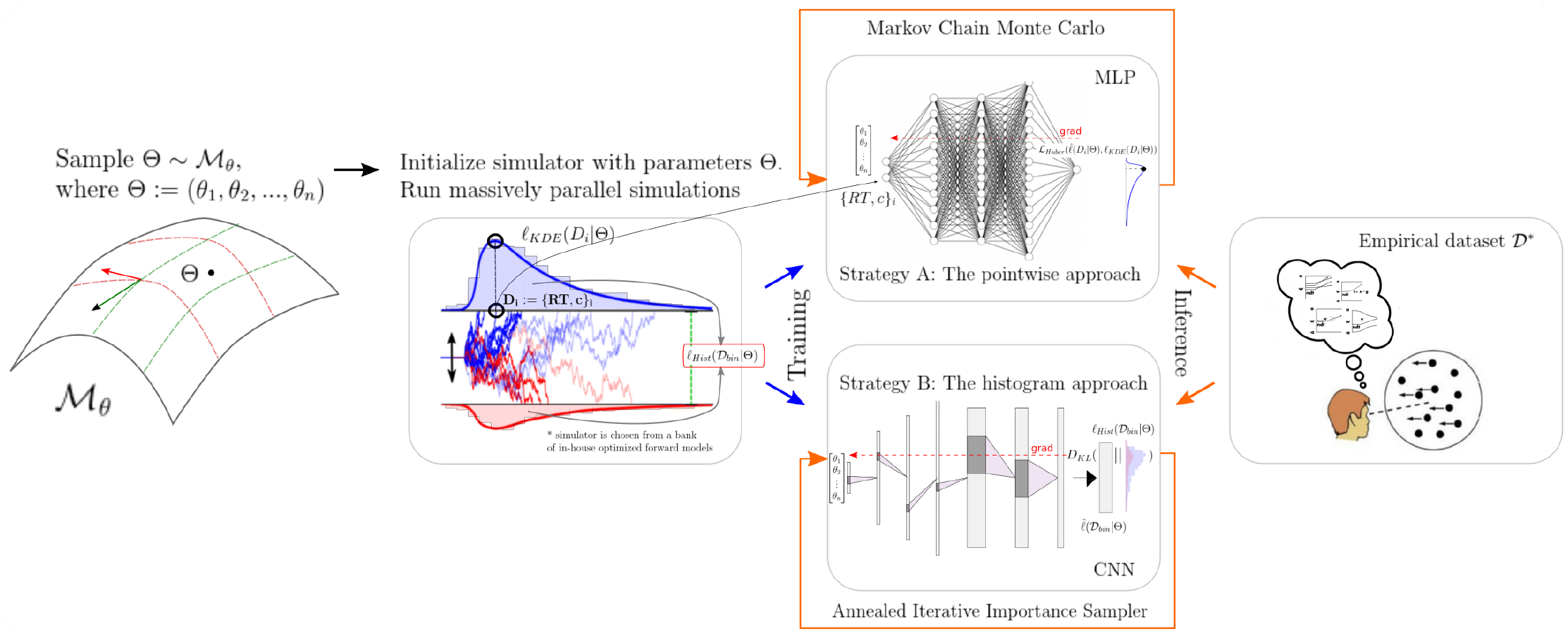
High level overview of our approaches. For a given model *M*, we sample model parameters *θ* from a region of interest (left 1), and run 100*k* simulations (left 2). We use those simulations to construct a KDE-based empirical likelihood, and a discretized (histogram-like) empirical likelihood. The combination of parameters and the respective likelihoods is then used to train the likelihood networks (right 1). Once trained we can use the MLP and CNN for posterior inference given an empirical / experimental dataset (right 2).

### Pointwise approach: Learn likelihoods of individual observations with MLPs

As a first approach, we use simple Multi-layered Perceptrons (MLPs), to learn the likelihood function of a given stochastic simulator. The network learns the mapping 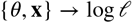, where *θ* represents the parameters of our stochastic simulator, **x** are the simulator outcomes (in our specific examples below, **x**, refers to the tuple (*rt, c*) of reaction times and choices), and 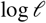 is the log-likelihood of the respective outcome. The trained network then serves as the approximate likelihood function 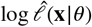. We emphasize that the network is trained as a function approximator, to provide us with a computationally cheap likelihood estimate, not to provide a surrogate simulator. Biological plausibility of the network is therefore not a concern when it comes to architecture selection.

### Histogram approach: Learn likelihoods of entire dataset distributions with CNNs

Our second approach is based on a convolutional neural network (CNN) architecture. Whereas the MLP learned to output a single scalar likelihood output for each data point (“trial”, given a choice, reaction time and parameterization), the goal of the CNN was to evaluate, for a given parameterization, the log-likelihood of an arbitrary number of datapoints via one forward pass through the network. To do so, the output of the CNN was trained to produce a probability distribution over a discretized version of the dataspace for a given parameterization of the stochastic model.

## 4 Test Beds

We choose variations of sequential sampling models (SSMs) common in the cognitive neurosciences as our test-bed (Figure 3). The range of models we consider allows for great flexibility in allowable data distributions (choices and response times). We believe that initial applications are most promising for such SSMs, because (i) analytic likelihoods are available for the most common variants (and thus provide an upper-bound benchmark for parameter recovery), and (ii) there exist many other interesting variants for which no analytic solution exists. Our methods however are quite general and any model that generates discrete choices and response times from which simulations can be drawn within a reasonable amount of time can be suitable to the amortization techniques discussed in this paper.

**Figure 3.**
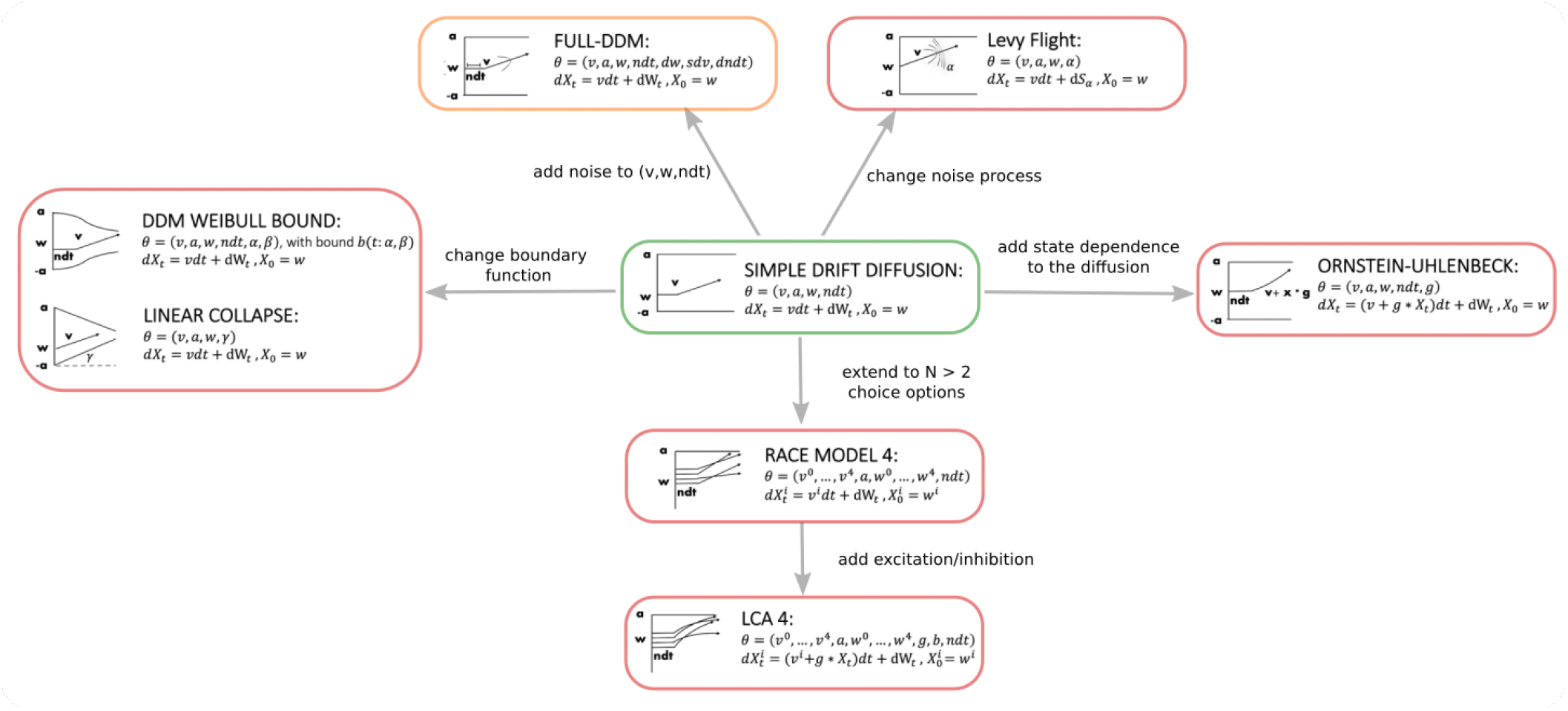
Pictorial representation of the stochastic simulators that form our test-bed. Our point of departure is the standard drift diffusion model (shown in green) due its analytical tractability and its prevalence as the most common SSM in cognitive neuroscience. By systematically varying different facets of the DDM, we test our LANs across a range of SSMs for parameter recovery, goodness of fit (posterior preditive checks) and inference runtime. While the FULL-DDM has an analytically tractable likelihood function that is expensive to compute (shown in orange), the other models do not possess a closed-form likelihood function (shown in red).

As a general principle, all models tested below are based on stochastic differential equations of the following form,

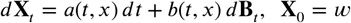

where we are concerned with the probabilistic behavior of the particle (or vector of particles) **x**. The behavior of this particle is driven by *a*(*t, x*), an underlying drift function, *b*(*t, x*) an underlying noise transformation function, *B*_*t*_ an incremental noise process, and *X*_0_ = *w* a starting point.

Of interest to us are specifically the properties of the first-passage-time-distributions (FPTD) for such processes, which are needed to compute the likelihood of a given response time/choice pair {*rt, c*}. In these models, the exit region of the particle (i.e., the specific boundary it reaches) determines the choice, and the time point of that exit determines the response time. The joint distribution of choices and reaction times are referred to as a FPTDs.

Given some exit-region *ԑ*, such FPTD’s are formally defined as,

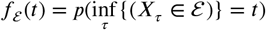

In words, a first passage time, for a given exit region, is defined as the first time point of entry into the exit region, and the FPTD is the probability, respectively of such an exit happening at any specified time *t*(***Ratcliff, 1978***; ***Feller, 1968***). Partitioning the exit region into subsets *ԑ*_1_, … *ԑ*_*n*_ (for example representing *n* choices), we can now define the set of defective distributions,

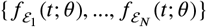

where *θ* ∈ Θ describes the collection of parameters driving the process. For every *ԑ*_*i’*_,

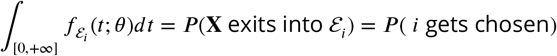

{*f*_≧_ … *f*_*ԑ*_} jointly define the FPTD such that,

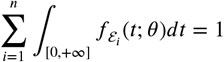

These functions 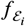, jointly serve as the likelihood function s.t.,

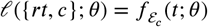

For illustration we focus the general model formulation above to the standard DDM. Details regarding the other models in our test-bed are relegated to the methods section at the end of the paper.

To obtain the DDM from the general equation above, we set *a*(*t, x*) = *v* (a fixed drift across time), *b*(*t, x*) = 1 (a fixed noise variance across time), and 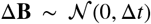. The DDM applies to the two alternative decision case, where decision correspond to to particle crossings of an upper or lower fixed boundary. Hence 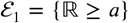 and 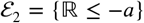 where *a* is a parameter of the model. The DDM also includes a normalized starting point *w* (capturing potential response biases or priors), and finally a non-decision time *τ* (capturing the time for perceptual encoding and motor output). Hence, the parameter vector for the DDM is then *θ* = (*v, a, w, τ*). The SDE is defined as,

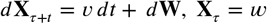

The DDM serves principally as a basic proof of concept for us, in that it is a model for which we can compute the exact likelihoods analytically (***Feller (1968)***;***Navarro and Fuss (2009)***).

The other models chosen for our test-bed systematically relax some of the fixed assumptions of the basic DDM, as illustrated in Figure 3.

We note that many new models can be constructed from the components tested here. As an example of this modularity, we introduce inhibition/excitation to the race model, which gives us the Leaky Competing Accumulator (LCA) (***Usher and McClelland, 2001***). We could then further extend this model by introducing parameterized bounds. We could introduce reinforcement learning parameters to a DDM (***Pedersen and Frank, 2020***) or in combination with any of the other decision models. Again we emphasize that while these diffusion-based models provide a large test-bed for our proposed methods, applications are in no way restricted to this class of models.

## 5 Results

### Networks learn likelihood function manifolds

Across epochs of training, both training and validation loss decrease rapidly and remain low (Figure 4A). Thus, over-fitting is not an issue, which is sensible in this context given that networks are given access to noiseless inputs and outputs and only have to learn a nearly deterministic transformation. This transformation is deterministic in the limit where the number of simulations on each sample tends to infinity. The low validation loss further shows that the network can interpolate likelihoods to specific parameter values it has not been exposed to (with the caveat that it has to be exposed to the same range; no claims are made about extrapolation).

**Figure 4.**
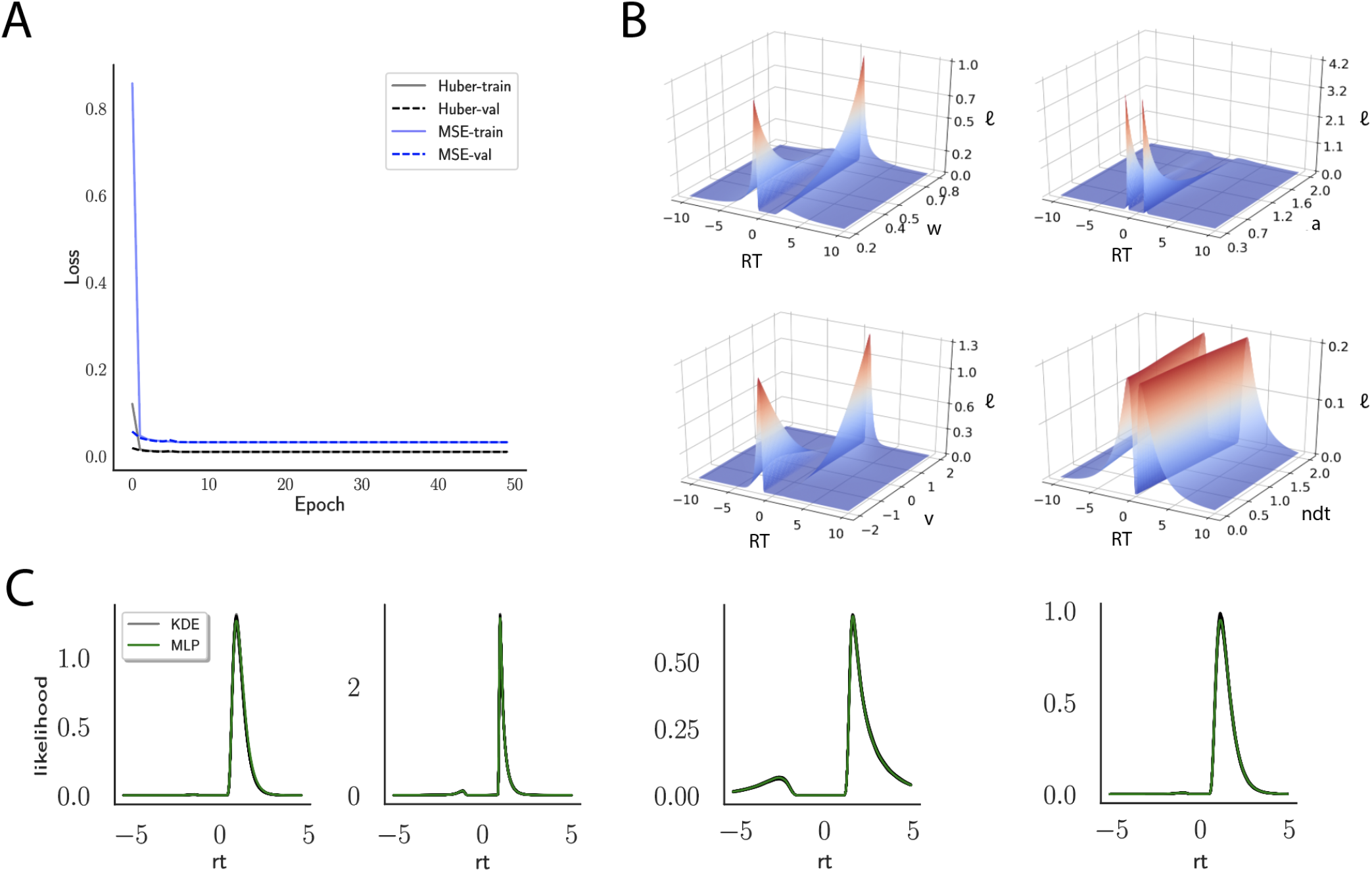
A Shows the training and validation loss for Huber as well as MSE for the DDM model. Training was driven by the Huber loss. **B** Illustrates the marginal likelihood manifolds for choices and RTs, by varying one parameter in the trained region. **C** shows MLP likelihoods in green for random model parameterizations, on top of a sample of 100 KDE-based empirical likelihoods derived from 100k samples each. The MLP mirrors the KDE likelihoods despite not having been explicitly trained on these parametrizations. Moreover, the MLP likelihood sits firmly at the mean of sample of 100 KDE’s

Indeed, a simple interrogation of the learned likelihood manifolds shows that they smoothly vary in an interpretable fashion with respect to changes in generative model parameters (Figure 4B). Moreover, Figure 4C shows that the MLP-likelihoods mirror those obtained by KDEs using 100,000 simulations, even though the model parameterizations were drawn randomly and thus not trained *per se*. We also note that the MLP-likelihoods appropriately filter out simulation noise (random fluctuations in the KDE empirical likelihoods across separate simulation runs of 100*K* samples each). This observation can also be gleaned from Figure 4C, which shows the learned likelihood to sit right at the center of sampled KDEs (note that for each subplot 100 such KDEs were used). As illustrated in the appendix, these observations hold across all tested models. One perspective on this is to consider the MLP-likelihoods as equivalent to KDE likelihoods derived from a much larger number of underlying samples and interpolated. The results for the CNN (not shown to avoid redundancy) mirror the MLP results. Finally, while Figure 4 depicts the learned likelihood for the simple DDM for illustration purposes, the same conclusions apply to the learned manifolds for all of the tested models (as shown in the appendix Figures 14–18). Indeed inspection of those manifolds is insightful for facilitating interpretation of the dynamics of the underlying models, how they differ from each other, and the corresponding RT distributions that can be captured.

### Parameter Recovery

#### Benchmark: Analytical Likelihood Available

While the above inspection of the learned manifolds is promising, a true test of the method is to determine whether one can perform proper inference of generative model parameters using the MLP and CNN. Such parameter recovery exercises are typically performed to determine whether a given model is identifiable for a given experimental setting (e.g., number of trials, conditions, etc). Indeed, when parameters are collinear, recovery can be imperfect even if the estimation method itself is flawless (***Wilson and Collins, 2019***; ***Nilsson et al., 2011***; ***Daw et al., 2011a***). A Bayesian estimation method, however, should properly assign uncertainty to parameter estimates in these circumstances, and hence it is also important to evaluate the posterior variances over model parameters.

Thus as a benchmark, we first consider the basic DDM for which an arbitrarily close approximation to the analytic likelihood is available (***Navarro and Fuss, 2009***). This benchmark allows us to compare parameter recovery given (1) the analytic likelihood, (2) an approximation to the likelihood specified by training an MLP on the analytical likelihood (thus evaluating the potential loss of information incurred by the MLP itself), (3) an approximation to the likelihood specified by training an MLP on KDE-based empirical likelihoods (thus evaluating any further loss incurred by the KDE reconstruction of likelihoods), and (4) an approximate likelihood resulting from training the CNN architecture. Figure 5 shows the results for the DDM.

**Figure 5.**
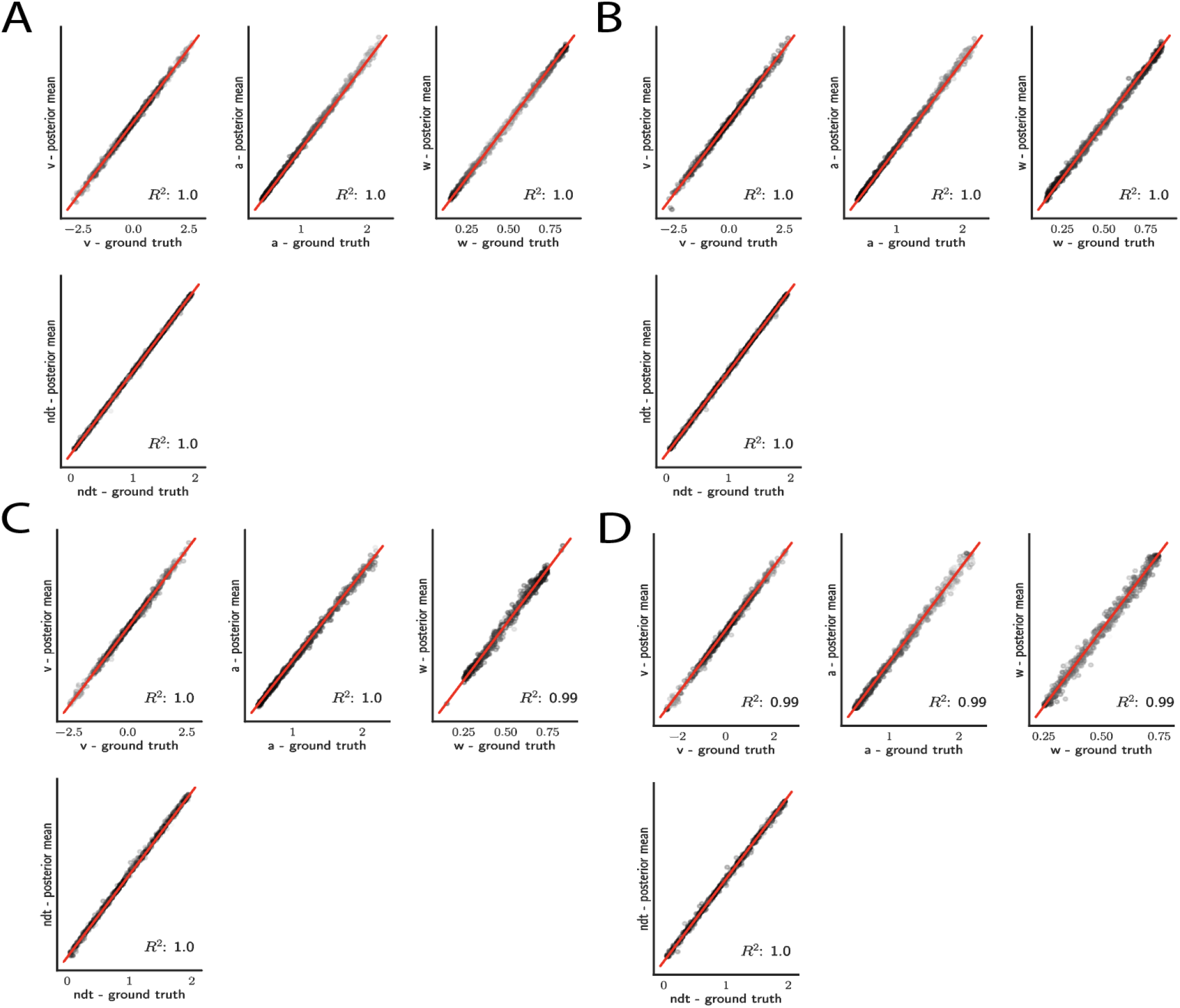
Simple DDM Parameter recovery results for, **A** analytic likelihood (ground truth), **B** MLP trained on analytic likelihood, **C** MLP trained on KDE-based likelihoods (100K simulations per KDE), **D** CNN trained on binned likelihoods. The results represent posterior means, based on inference over datasets of size *N*_1_ = 1024 “trials”. Dot shading is based on parameter-wise normalized posterior variance, with lighter shades indicating larger posterior uncertainty of the parameter estimate.

For the simple DDM and analytic likelihood, parameters are nearly perfectly recovered given *N* = 1024 data points (“trials”) (Figure 5A). Notably, these results are mirrored when recovery is performed using the MLP trained on the analytic likelihood (Figure 5B). This finding corroborates, as visually suggested by the learned likelihood manifolds, the conclusion that globally the likelihood function was well behaved. Moreover, only slight reductions in recoverability were incurred when the MLP was trained on the KDE likelihood estimates (Figure 5C), likely due to the known small biases in KDE itself (***Turner et al., 2015b***). Similar performance is achieved using the CNN instead of MLP (Figure 5 D).

As noted above, an advantage of Bayesian estimation is that we obtain an estimate of the posterior uncertainty in estimated parameters. Thus, a more stringent requirement is to additionally recover the correct posterior variance, for a given dataset 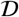 and model 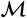. One can already see visually in Figure 5C, D that posterior uncertainty is larger when the mean is further from the ground truth (lighter shades of grey indicate higher posterior variance). However to be more rigorous one can assess whether the posterior variance is precisely what it should be.

The availability of an analytical likelihood for the DDM, together with our use of sampling methods (as opposed to variational methods which can severely bias posterior variance), allows us to obtain the “ground truth” uncertainty in parameter estimates. Figure 6 shows that the sampling from a MLP trained on analytical likelihoods, a MLP trained on KDE-based likelihoods and a CNN all yield excellent recovery of the variance. For an additional run that involved datasets of size *n* = 4096 instead of *n* = 1024, we observed a consistent decrease in posterior variance across all methods (not shown) as expected.

**Figure 6.**
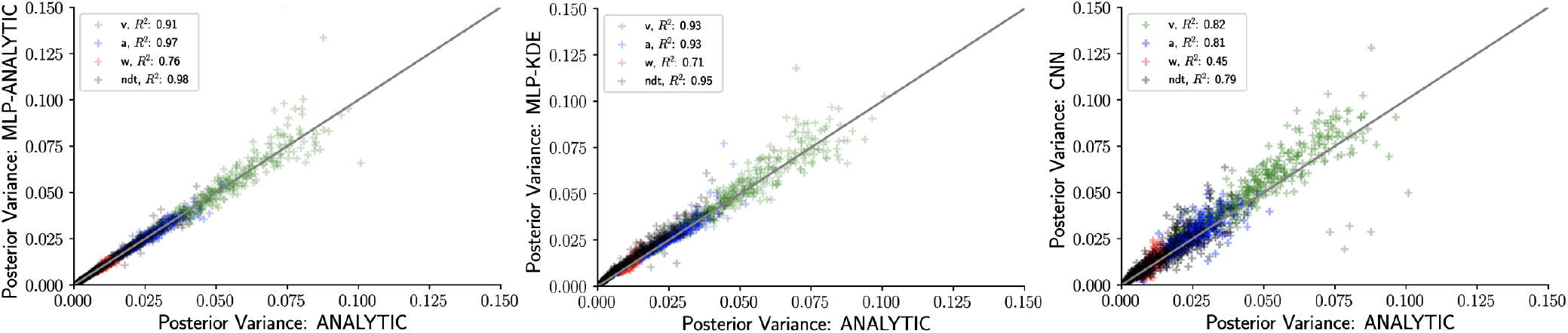
Inference using LANs recovers posterior uncertainty. Here we leverage the analytic solution for the DDM to plot the “ground truth” posterior variance on the x-axis, against the posterior variance from the LANs on the y-axis. Left. MLPs trained on the analytic likelihood. Middle. MLPs trained on KDE-based emprical likelihoods. Right. CNNs trained on binned empirical likelihoods. Datasets were equivalent across methods for each model (left to right) and involved *n* = 1024 samples.

#### No Analytical Likelihood Available

As a proof of concept for the more general ABC setting, we show parameter recovery results for two non-standard models, the ANGLE and WEIBULL models as described in the the Test Bed section. The results are summarized in Figure 7 and 8 and described in more detail in the following two paragraphs.

**Figure 7.**
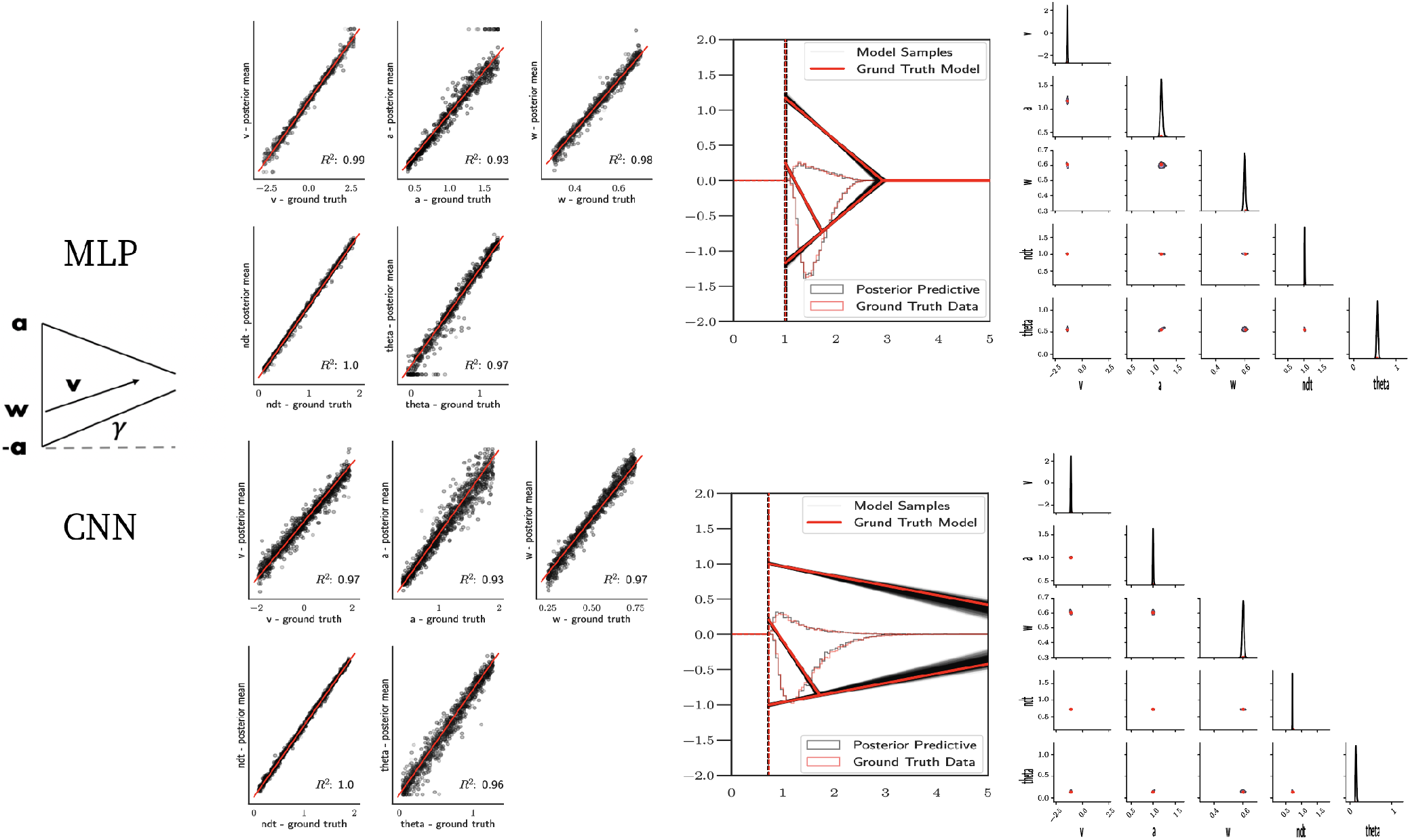
ANGLE model parameter recovery and posterior predictives. **Left.** Parameter recovery results for the MLP (top) and textitCNN (bottom). **Right.** Posterior predictive plots for two representative datasets. Model samples of all parameters (black) match those from the true generative model (red), but one can see that for the lower dataset, the bound trajectory is somewhat more uncertain (more dispersion of the angle samples). In both cases, the posterior predictive (black histograms) is shown as predicted choice proportions and RT distributions for upper and lower boundary responses, overlaid on top of the ground truth data (red; hardly visible since overlapping / matching).

**Figure 8.**
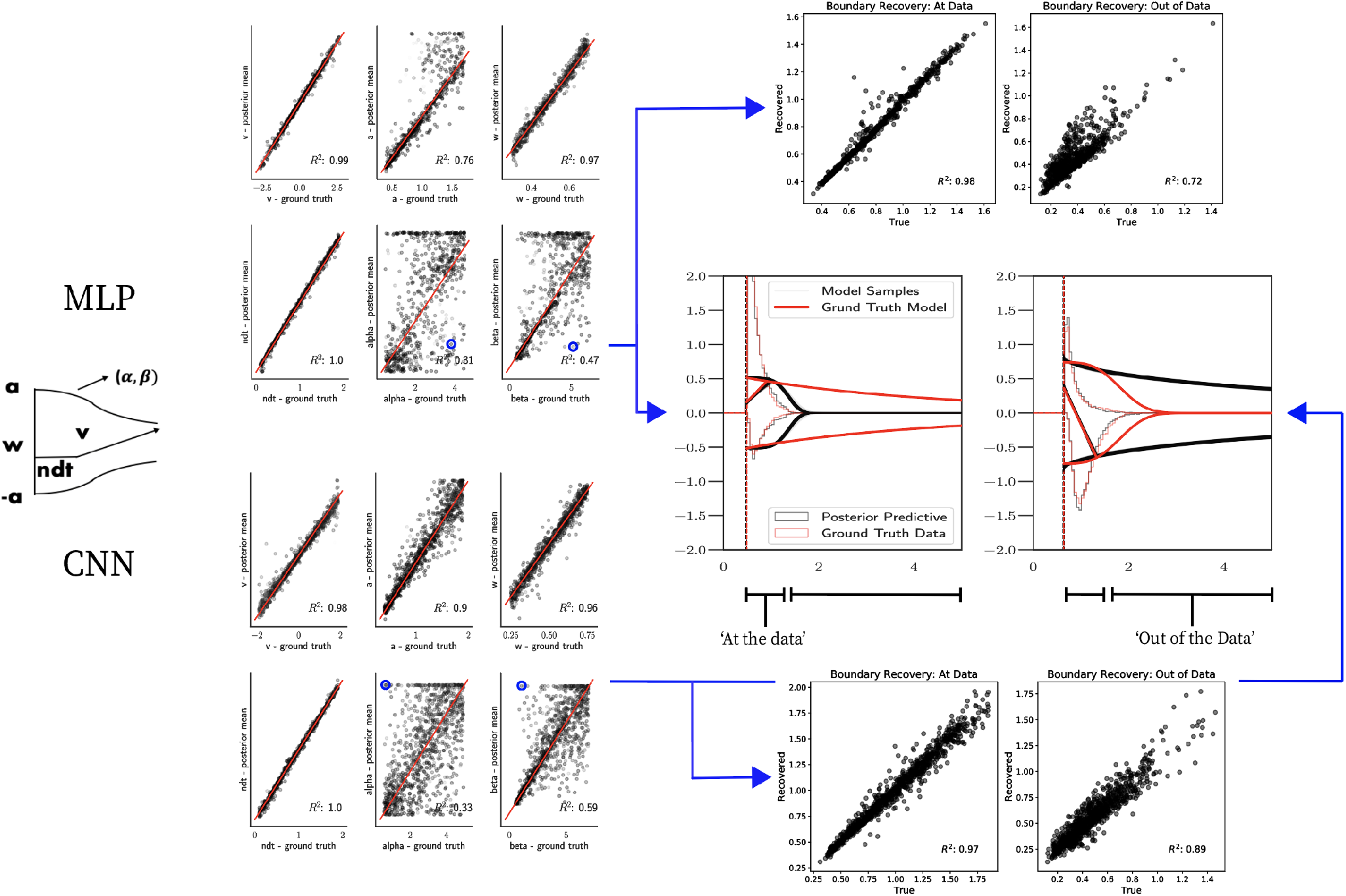
WEIBULL model parameter recovery and posterior predictives.**Left.** Parameter recovery results for the MLP (top) and CNN (bottom). **Right.** Posterior predictive plots for two representative datasets in which parameters were poorly estimated (denoted in blue on the left). In these examples, model samples (black) recapitulate the generative parameters (red) for the non-boundary parameters, the recovered bound trajectory is poorly estimated relative to the ground truth, despite excellent posterior predictives in both cases (RT distributions for upper and lower boundary, same scheme as Figure 7). Nevertheless, one can see that the net decision boundary is adequately recovered within the range of the RT data that are observed. Across all datasets, the net boundary 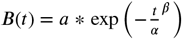 is well recovered within the range of the data observed, and somewhat less so outside of the data, despite poor recovery of individual Weibull parameters *α* and *β*.

##### Parameter Recovery

Figure 7 shows that both the MLP and CNN methods consistently yield very good to excellent parameter recovery performance for the ANGLE model, with parameter-wise regression coefficients globally above *R*^2^ > 0.9. As shown in Figure 8 parameter recovery for the WEIBULL model, is less successful however, particularly for the weibull collapsing bound parameters. The drift parameter *v*, the starting point bias *w* and the non-decision time are estimated well, however the boundary parameters *a*, *α* and *β* are less well recovered by the posterior mean. Judging by the parameter recovery plot, the MLP seems to perform slightly less well on the boundary parameters when compared to the CNN.

To interrogate the source of the poor recovery of *α* and *β* parameters, we considered the possibility that the model itself may have issues with identifiability, rather than poor fit. Figure 8 shows that indeed, for two representative datasets in which these parameters are poorly recovered, the model nearly perfectly reproduces the ground truth data in the posterior predictive RT distributions. Moreover, we find that whereas the individual Weibull parameters are poorly recovered, the net boundary *B*(*t*) is very well recovered, particularly when evaluated within the range of the observed dataset. This result is reminiscent of the literature on sloppy models (***Gutenkunst et al., 2007***), where sloppiness implies that various parameter configurations can have the same impact on the data. Moreover, two further conclusions can be drawn from this analysis. First, when fitting the WEIBULL model, researchers should interpret the bound trajectory as a latent parameter rather than the individual *α* and *β* parameters *per se*. Second, the WEIBULL model may be considered as viable only if the estimated bound trajectory varies sufficiently within the range of the empirical RT distributions. If the bound is instead flat or linearly declining in that range, the simple DDM or ANGLE models may be preferred, and their simpler form would imply that they would be selected by any reasonable model comparison metric.

Finally, the appendix shows parameter recovery studies on a number of other stochastic simulators with non-analytic likelihoods, described in the Test Bed section. Appendix 0 shows a table of parameter wise recovery *R*^2^ for all models tested. In general, recovery ranges from good to excellent. Given the WEIBULL results above, we attribute the less good recovery for some of these models to identifiability issues and specific dataset properties rather than to the method *per se*. We note that our parameter recovery studies here are in general constrained to the simplest inference setting equivalent to a single subject, single condition experimental design. Moreover we use uninformative priors for all parameters of all models. Thus these results provide a lower bound on parameter recoverability; please see Section 6 for how recovery can benefit from more complex experimental designs with additional task conditions, which more faithfully represents the typical inference scenario deployed by cognitive neuroscientists.

### Runtime

A major motivation for this work is the amortization of network training time during inference, affording researchers the ability to test a variety of theoretically interesting models for linking brain and behavior without large computational cost. To quantify this advantage we provide some results on the posterior sampling run-times using (1) the MLP with Slice Sampling (***Neal, 2003***) and (2) CNN with iterated importance sampling.

The MLP timings are based on slice sampling (***Neal, 2003***), with a minimum of *n* = 2000 samples. The sampler was stopped at some *n* > = 2000, for which the Geweke statistic (***Geweke, 1992***) indicated convergence (the statistic was computed once every 100 samples for *n* > = 2000). Using an alternative sampler, based on DEMCMC and stopped when the Gelman Rubin 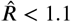 (***Gelman et al., 1992***) yielded very similar timing results and was omitted in our figures.

For the reported importance sampling runs we used 200*K* importance samples per iteration, starting with *γ* values of 64, which was first reduced to 1 where in iteration 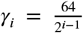, before a stopping criterion based on relative improvement of the confusion metric was used.

Figure 9**A** shows that all models can be estimated in the order of hundreds of seconds (minutes), comprising a speed improvement of at least two orders of magnitude compared to traditional ABC methods using KDE during inference (i.e., the PDA method motivating this work (***Turner et al., 2015b***)). Indeed, this estimate is a lower bound on the speed-improvement: we extrapolate only the observed difference between network evaluation and online simulations, ignoring the additional cost of constructing and evaluating the KDE-based likelihood. We decided to use this benchmark because it provides a fairer comparison to more recent PDA approaches in which the KDE evaluations can be sped up considerably ***Holmes (2015)***.

**Figure 9.**
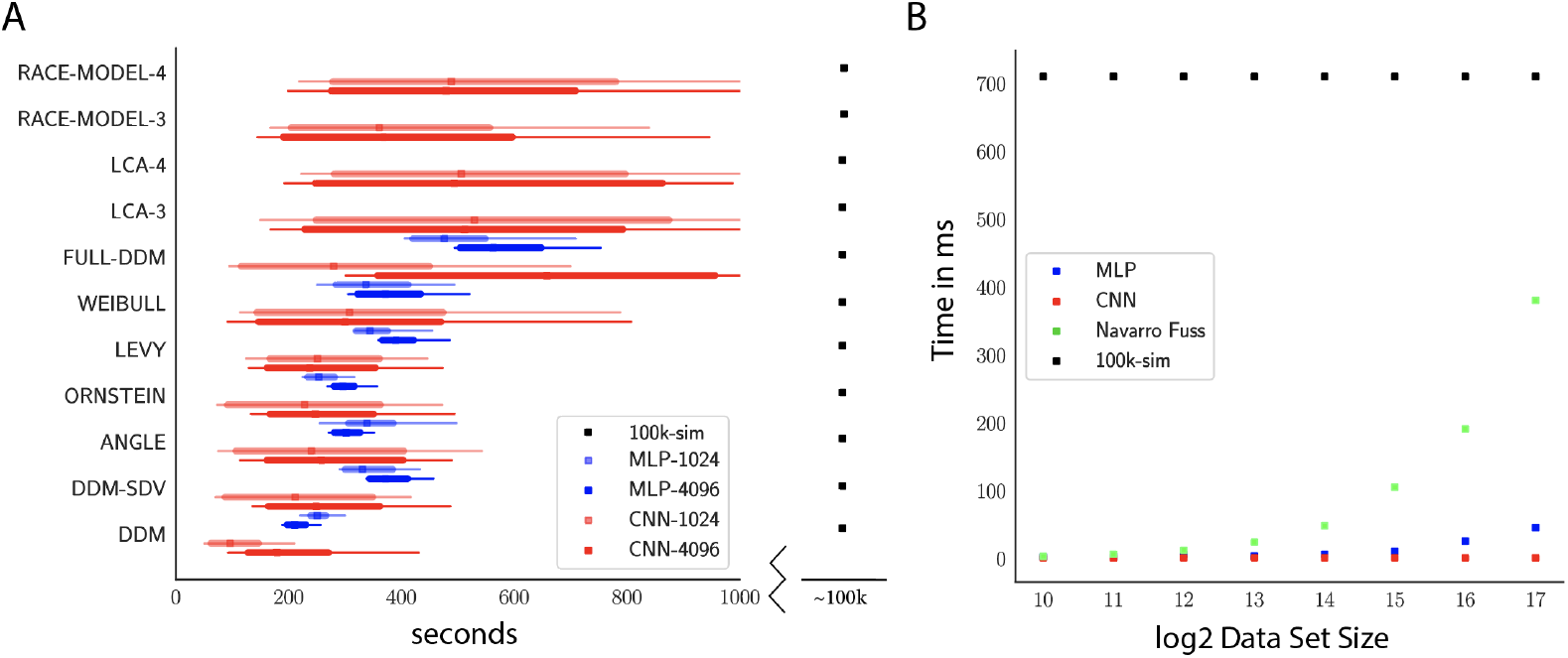
A Comparison of sampler timings for the MLP and CNN methods, for datasets of size 1024 and 4096. For comparison, we include a lower bound estimate of the sample timings using traditional PDA approach during online inference (using 100k online simulations). (100K simulations were used because we found this to be required for sufficiently smooth likelihood evaluations and is the number of simulations used to train our networks; fewer samples can of course be used at the cost of worse estimation, and only marginal speed up). Note further that these timings scale with the number of participants and task conditions for the online method, but not for LANs, where they can be parallelized. **B** compares the timings for obtaining a single likelihood evaluation for a given dataset. MLP and CNN refer to Tensorflow implementations of the corresponding networks. Navarro Fuss, refers to a cython (cpu) implementation of the algorithm suggested ***Navarro and Fuss (2009)*** for fast evaluation of the analytical likelihood of the DDM. 100k-sim refers to the time it took a highly optimized cython (cpu) version of a DDM-sampler to generate 100k simulations (averaged across 100 parameterizations).

Notably, due to its potential for parallelization (especially on GPUs), our neural network methods can even induce performance speed-ups relative to analytic likelihood evaluations. Indeed, figure 9**B** shows that as the dataset grows, runtime is significantly faster than even a highly optimized cython implementation of the Navarro Fuss algorithm (***Navarro and Fuss, 2009***) algorithm for evaluation of the analytic DDM likelihood. This is also noteworthy in light of the Full-DDM model (as described in the test-bed section), for which it is currently common to compute the likelihood term via quadrature methods, in turn based on repeated evaluations of the Navarro Fuss algorithm, which can easily inflate the evaluation time by 1 to 2 orders of magnitude. In contrast, evaluation times for the MLP and CNN are only marginally slower (as a function of the slightly larger network size in response to higher dimensional inputs). We confirm (omitted as separate Figure) from experiments with the HDDM python toolbox, that our methods end up approximately 10 − 50 times faster for the Full-DDM than the current implementation based on numerical integration, maintaining comparable parameter recovery performance. We strongly suspect there to be additional remaining potential for performance optimization.

## 6 Hierarchical inference

One of the principal benefits of LANs is that they can be directly extended – without further training – to arbitrary hierarchical inference scenarios, including those in which (i) individual participant parameters are drawn from group distributions; (ii) some parameters are pooled and others separated across task conditions and (iii) neural measures are estimated as regressors on model parameters (Figure 10). Hierarchical inference is critical for improving parameter estimation particularly for realistic cognitive neuroscience datasets in which thousands of trials are not available for each participant, and/or where one estimates impacts of noisy physiological signals onto model parameters (***Wiecki et al., 2013***; ***Boehm et al., 2018***; ***Vandekerckhove et al., 2011***; Ratcliff and Childers, ????).

**Figure 10.**
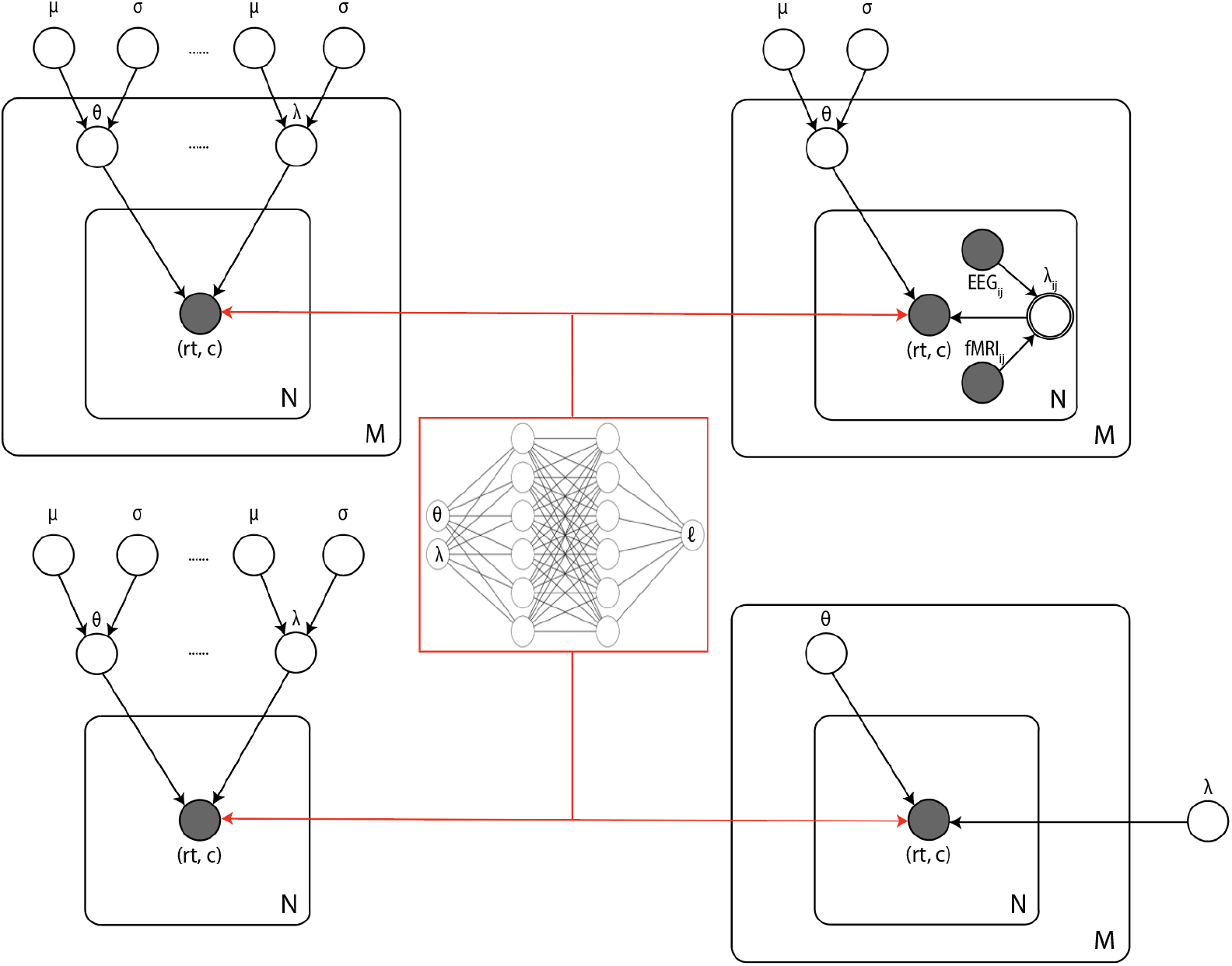
Illustrates common inference scenarios applied in the cognitive neurosciences and enabled by our amortization methods. (Upper left) basic hierarchical model across **M** participants, with **N** observations (trials) per participant. Parameters for individuals are assumed to be drawn from group distributions. (Upper right) hierarchical models which further estimate the impact of trial-wise neural regressors onto model parameters. (Lower left) non-hierarchical, standard model estimating one set of parameters across all trials. (Lower right), common inference scenario in which a subset of parameters (*θ*) are estimated to vary across conditions **M**, while others (*λ*) are global. LANs can be immediately re-purposed for all of these scenarios (and more) without further training.

To provide a proof of concept, we developed an extension to the HDDM python toolbox ***Wiecki et al. (2013)***, widely used for hierarchical inference of the DDM applied to such settings. Lifting the restriction of previous versions of HDDM to only DDM variants with analytical likelihoods, we imported the MLP likelihoods for all two-choice models considered in this paper. Note that GPU-based computation is supported out of the box, which can easily be exploited with minimal over-head using free versions of Google’s Colab notebooks. We generally observed GPUs to improve speed approximately five-fold over CPU-based setups for the inference scenarios we tested.^2^

Figure 11 shows example results from hierarchical inference using the ANGLE model, applied to synthetic datasets comprising 5 and 20 subjects (a superset of participants). Recovery of individual parameters was adequate even for 5 participants, and we also observe the expected improvement of recovery of the group level parameters *μ* and *σ* for 20 participants.

**Figure 11.**
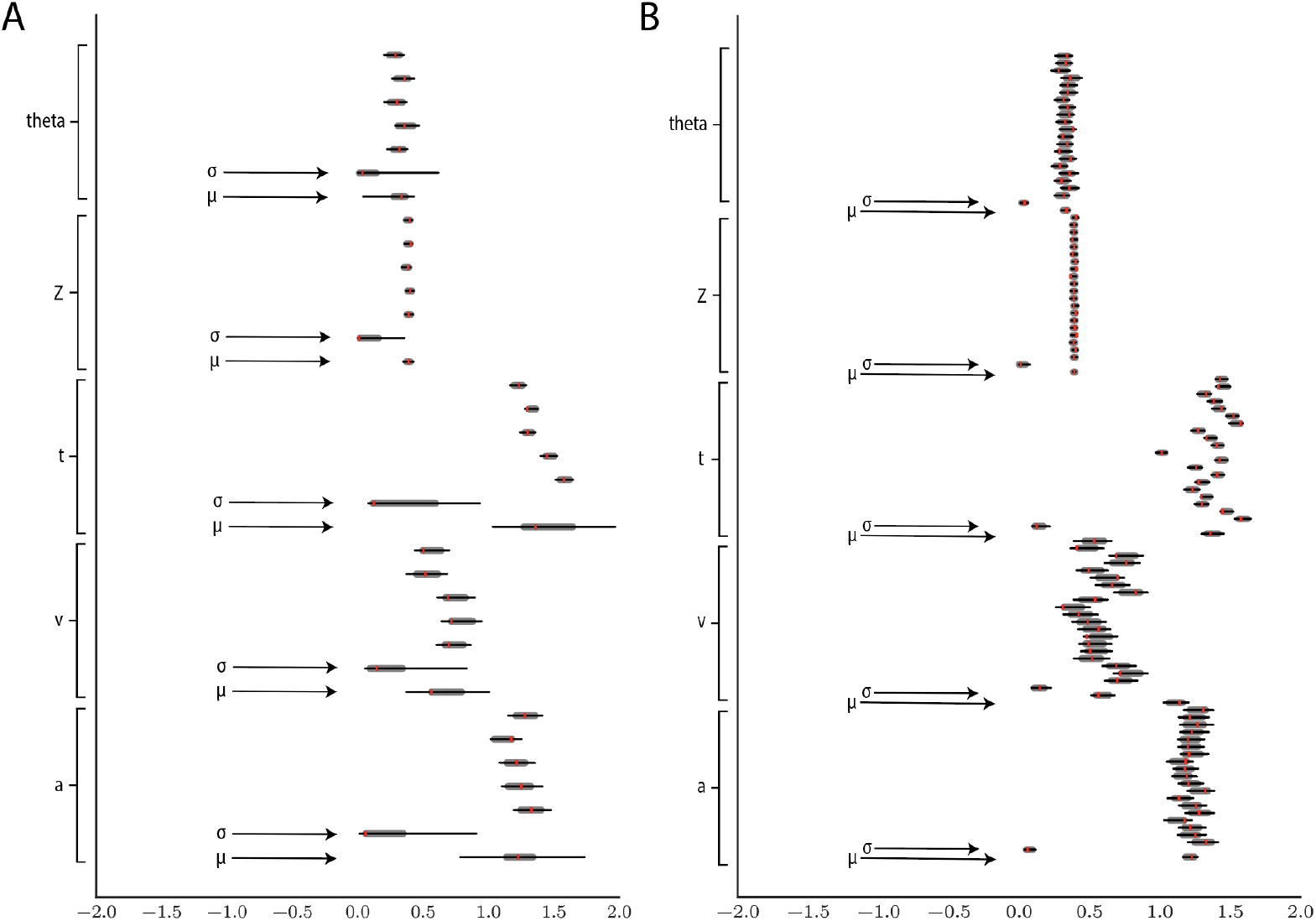
Hierarchical inference results using the MLP likelihood imported into the HDDM package. **A** posterior inference for the ANGLE model on a synthetic dataset with 5 participants and 500 trials each. Posterior distributions are shown with caterpillar plots (thick lines correspond to 5 − 95 percentiles, thin lines correspond to 1 − 99 percentiles) grouped by parameters (ordered from above {*subject*_1_, …, *subject*_*n*_, *μ*_*group*_*σ*_*group*_}). Ground truth simulated values denoted in red. **B** hierarchical inference for synthetic data comprising 20 participants and 500 trials each. *μ* and *σ* indicate the group level mean and variance parameters. Estimates of group level posteriors improve with more participants as expected with hierarchical methods. Individual level parameters are highly accurate for each participant in both scenarios.

Figure 12 shows an example that illustrates how parameter recovery is affected when a dataset contains multiple experimental conditions (e.g., different difficulty levels). It is common in such scenarios to allow task conditions to affect a single (or subset) of model parameters (in the cases shown: *v*), while other model parameters are pooled across conditions. As expected, for both the Full-DDM (panel **A**) and the Levy model (panel **B**), the estimation of global parameters is improved when increasing the number of conditions from 1 to 5 to 10 (left to right, where the former are subsets of the latter datasets. These experiments confirm that one can more confidently estimate parameters that are otherwise difficult to estimate, such as the noise *α* in the Levy model and *sv* the standard deviation of the drift in the Full-DDM.

**Figure 12.**
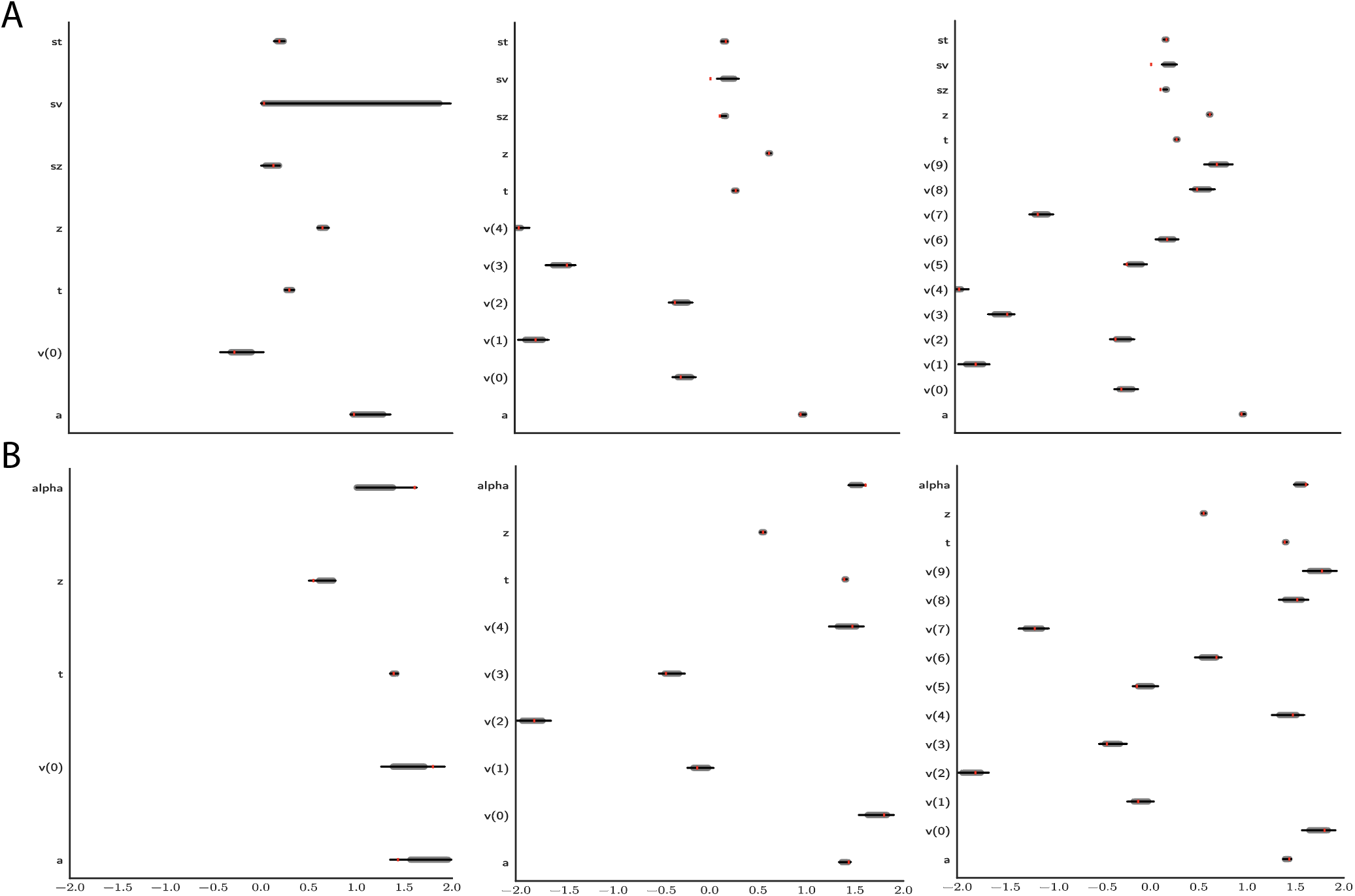
Effect of multiple experimental conditions on inference. The panel shows an example of posterior inference for **1**, (left), **5**(middle) and **10**(right) conditions. **A** and **B**, refer to the Full-DDM and Levy models respectively. The drift parameter *v* is estimated to vary across conditions, while the other parameters are treated as global across conditions. Inference tends to improve for all global parameters when adding experimental conditions. Importantly this is particularly evident for parameters that are otherwise notoriously difficult to estimate, such as *sv* (trial by trial variance in drift in the Full-DDM model) and *α* (the noise distribution in the Levy model). Red stripes show the ground truth values of the given parameters.

Both of these experiments provide evidence that our MLPs provide approximate likelihoods which behave in accordance with what is expected from proper analytic methods, while also demonstrating their robustness to other samplers (i.e., we used HDDM slice samplers without further modification for all models).

We expect that proper setting of prior distributions (uniform in our examples) and further refinements to the slice sampler settings (to help mode discovery), can improve these results even further. We include only the MLP method in this section, since it is most immediately amenable to the kind of trial-by-trial level analysis that HDDM is designed for. We plan to investigate the feasibility of including the CNN method into HDDM in future projects.

## 7 Discussion

Our results demonstrate the promise and potential of amortized likelihood approximation networks for Bayesian parameter estimation of neurocognitive process models. Learned manifolds and parameter recovery experiments showed successful inference using a range of network architectures and posterior sampling algorithms, demonstrating the robustness of the approach.

Although these methods are extendable to any model of similar complexity, we focused here on a class of sequential sampling models, primarily because the most popular of them – the DDM – has an analytic solution, and is often applied to neural and cognitive data. Even slight departures from the standard DDM framework (e.g., dynamic bounds, or changes in the noise distribution) are often not considered for full Bayesian inference due to the computational complexity associated with traditional ABC methods. We provide access to the learned likelihood functions (in the form of network weights) and code to enable users to fit a variety of such models with orders of magnitude speed up (minutes instead of days). In particular, we provided an extension to the commonly used HDDM toolbox (***Wiecki et al., 2013***) that allows users to apply these models to their own datasets immediately. We also provide access to code that would allow users to train their own likelihood networks and perform recovery experiments, which can then be made available to the community. We offered two separate approaches with their own relative advantages and weaknesses. The MLP is suited for evaluating likelihoods of individual observations (choices, response times) given model parameters, and as such can be easily extended to hierarchical inference settings and trial-by-trial regression of neural activity onto model parameters. We showed that importing the MLP likelihood functions into the HDDM toolbox affords fast inference over a variety of models without tractable likelihood functions. Moreover, these experiments demonstrated that use of the neural network likelihoods even confers a performance speed up over the analytic likelihood function – particularly for the Full-DDM, which otherwise required numerical methods on top of the analytic likelihood function for the simple DDM.

Conversely, the CNN approach is well suited for estimating likelihoods across parameters for entire datasets in parallel, as implemented with importance sampling. More generally and implying potential further improvements, any sequential monte carlo (SMC) method may be applied instead. These methods offer a more robust path to sampling from multimodal posteriors compared to MCMC, at the cost of the curse of dimensionality, rendering them potentially less useful for highly parameterized problems, such as those that require hierarchical inference. Moreover, representing the problem directly as one of learning probability distributions, and enforcing the appropriate constraints by design endows the CNN approach with a certain conceptual advantage. Finally we note that in principle (with further improvements) trial-level inference is possible with the CNN approach, and vice-versa, importance sampling can be applied to the MLP approach.

In this work, we employed sampling methods (MCMC and importance sampling) for posterior inference, because in the limit they are well known to allow for accurate estimation of posterior distributions on model parameters, including not only mean estimates but their variances and covariances. Accurate estimation of posterior variances is critical for any hypothesis testing scenario, because it allows one to be confident about the degree of uncertainty in parameter estimates. Indeed, we showed that for the simple DDM, we found that posterior inference using our networks yielded nearly perfect estimation of the variances of model parameters (which are available due to the analytic solution). Of course, our networks can also be deployed for other estimation methods even more rapidly: they can be immediately used for maximum likelihood estimation via gradient descent, or within other approximate inference methods, such as variational inference (see ***Acerbi (2020)*** for a related approach).

Other approaches exist for estimating generalized diffusion models. A recent example, not discussed thus far, is the pyDDM Python toolbox (***Shinn et al., 2020***), which allows maximum likelihood estimation of generalized drift diffusion models (GDDM). The underlying solver is based on the Fokker-Planck equations, which allow access to approximate likelihoods (where the degree of approximation is traded off with computation time / discretization granularity) for a flexible class of diffusion-based models, notably allowing arbitrary evidence trajectories, starting point and non-decision time distributions. However, to incorporate trial-by-trial effects would severely inflate computation time (on the order of the number of trials), since the solver would have operate on a trial by trial level. Moreover, any model that is not driven by Gaussian Diffusion, such as the Levy model we considered here, or the linear ballistic accumulator, is out of scope with this method. In contrast LANs can be trained to estimate any such model, limited only by the identifiability of the generative model itself. Finally, pyDDM does not afford full Bayesian estimation and thus quantification of parameter uncertainty and covariance.

By focusing on likelihood approximation networks, our approach affords the flexibility of networks serving as plug-ins for hierarchical or arbitrarily complex model extensions. In particular, the networks can be immediately transferred, without further training, to arbitrary inference scenarios in which researchers may be interested in evaluating links between neural measures and model parameters, and to compare various assumptions about whether parameters are pooled and split across experimental manipulations. This flexibility in turn sets our methods apart from other amortization and neural network based ABC approaches offered in the statistics, machine learning and computational neuroscience literature (***Papamakarios and Murray, 2016***; ***Papamakarios et al., 2019a***; ***Goncalves et al., 2020***; ***Lueckmann et al., 2019***; ***Radev et al., 2020a, b***), while staying conceptually extremely simple. Instead of focusing on extremely fast inference for very specialized inference scenarios, our approach focuses on achieving speedy inference while not implicitly compromising modeling flexibility through amortization step.

## 8 Limitations and Future Work

There are several limitations of the methods presented in this article, which we hope to address in future work. While allowing for great flexibility, the MLP approach suffers from the drawback that we don’t enforce (or exploit) the constraint that 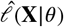 is a valid probability distribution, and hence the networks have to learn this constraint implicitly and approximately. Enforcing this constraint, has the potential to improve estimation of tail probabilities (a known issue for KDE approaches to ABC more generally (***Turner et al., 2015b)***).

The CNN encapsulation exploits the fact that 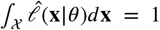, however makes estimation of trial-by-trial effects more resource hungry.

We plan to investigate the potential of the CNN for trial-by-trial estimation in future research.

A potential solution that combines the strengths of both the CNN and MLP methods, is to utilize Mixture Density Networks to encapsulate the likelihood functions. We are currently exploring this avenue. Mixture density networks have been successfully applied in the context of ABC (***Papamakarios and Murray, 2016***), however training can be unstable without extra care (***Guillaumes, 2017***). Similarly, invertible flows may be used to learn likelihood functions (***Papamakarios et al., 2019b***), however the philosophy remains focused on distributions of summary statistics for single datasets. While impressive improvements have materialized at the intersection of ABC and Deep Learning methods (***Papamakarios et al., 2019a***; ***Greenberg et al., 2019***; ***Goncalves et al., 2020***), generally less attention has been paid to amortization methods that are not only of case-specific efficiency but sufficiently modular to serve a large variety of inference scenarios (e.g., Figure 10). This is an important gap, which we believe the popularization of the powerful ABC framework in the domain of experimental science hinges upon. A second, and short-term avenue for future work is the incorporation of our presented methods into the HDDM Python Toolbox (***Wiecki et al., 2013***) to extend its capabilities to a larger variety of SSMs. Initial work in this direction is completed, alpha version of the extension being available in form of a tutorial under https://github.com/lnccbrown/al-nets/tree/master/hddmnn-tutorial.

Our current training pipeline can be further optimized on two fronts. First, no attempt was made to minimize the size of the network needed to reliably approximate likelihood functions so as to further improve computational speed. Second, little attempt was made to optimize the amount of training provided to networks. For the models explored here, we found it sufficient to simply train the networks for a very large number of simulated datapoints such that interpolation across the manifold was possible. However, as model complexity increases, it would be useful to obtain a measure of the networks’ uncertainty over likelihood estimates for any given model parameterization. Such uncertainty estimates would be beneficial for multiple reasons. One such benefit would be to provide a handle on the reliability of sampling, given the parameter region. Moreover, such uncertainty estimates could be used to guide the online generation of training data to train the networks in regions with high uncertainty. At the intersection of ABC and Neural Networks active learning has been explored via uncertainty estimates based on network ensembles (***Lueckmann et al., 2019***). We plan to additionally explore the use of Bayesian neural networks, which provide uncertainty over their weights, for this purpose (***Neal, 1995***).

One more general shortcoming of our methods is the reliance on empirical likelihoods for training, which in turn are based on a fixed number of samples across model parameterizations, just as the PDA method proposed by ***Turner et al. (2015b)***. Recently this approach has been criticized fundamentally on grounds of producing bias in the generated KDE based likelihood estimates (***van Opheusden et al., 2020***). A reduction of the approximate likelihood problem to one of inverse binomial sampling was proposed (***van Opheusden et al., 2020***), which will generate unbiased likelihood estimates. To address this concern, we will investigate adaptive strategies for the selection of the simulations count *n*. We however highlight two points here. First, our networks add interpolation to the actual estimation of a likelihood. Likelihoods close in parameter space therefore share information, which translates into an effectively higher simulation count than the 100*k* chosen to construct each empirical likelihood used for training. Quantifying this benefit precisely we leave for future research, however we suspect, as suggested by Figure 4, that it may be substantial. Second, while we generally acknowledge that bias in the tails remains somewhat of an issue, resolution is at best partial even in the proposed methods of (***van Opheusden et al., 2020***). For the estimation of parameterizations for which a given datapoint is extremely unlikely, the authors suggest to threshold the simulation count so that their algorithm is guaranteed to stop. This effectively amounts to explicitly allowing for bias again. Although this approach is elegant conceptually, excessive computation remains an issue in application, especially when we need accurate estimates of the tail-end of likelihood functions.

Furthermore we relegate to future research proper exploitation of the fact that LANs are by design differentiable in the parameters. We are currently working on an integration of LANs with tensorflow probability (***Abadi et al., 2016***), utilizing autograd to switch our MCMC method to the gradient-based NUTS sampler (***Matthew and Andrew, 2014***). Main benefits of this sampler are robust mixing behavior, tolerance for high levels of correlations in the parameter space, while at the same time maintaining the ability to sample from high dimensional posteriors. High level of correlations in posteriors are traditionally an achilles heel of the otherwise robust coordinate wise slice samplers. DEMCMC and Iterated Importance samplers are somewhat more robust in this regards, however both may not scale efficiently to high dimensional problems. Robustness concerns aside, initial numerical results additionally show some promising further speed-ups.

Lastly, in contrast to the importance sampler driving the posterior inference for the CNN, we believe that some of the performance deficiencies of the MLP, are the result of our Markov Chains not having converged to the target distribution. A common problem seems to be that the sampler hits the bounds of the constrained parameter space and does not recover from that. As we show in Figure 7 and Figure 8, even ostensibly bad parameter recoveries follow a conceptual coherence and lead to good posterior predictive performance. We therefore may be under-reporting the performance of the MLP and plan to test the method on an even more comprehensive suite of MCMC samplers, moreover including thus far neglected potential for re-parameterization.

## 9 Acknowledgements

We thank Michael Shvartsman, Matthew Nassar and Thomas Serre for helpful comments and discussion regarding earlier versions of this manuscript. Furthermore, we thank Mads Lund Pederson for help with integrating our methods into the HDDM python toolbox.

## 10 Additional Methods

### Test-Beds

#### General Information

All models were simulated using the Euler-Maruyama method, which for some fixed discretization step-size Δ*t*, evolves the process as,

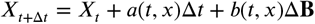

where the definition of Δ**B** depends on the noise process. For simple Brownian Motion this translates into gaussian displacements, specifically 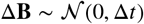, which is commonly denoted as *d***W**. More generally the noise need not be Gaussian, and indeed we later apply our methods to the Levy Flight model for which the noise process is an alpha stable distribution, denoted as **L**_*α*_ s.t. 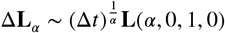.

The models chosen for our test-bed systematically vary different aspects of complexity, as illustrated in 3. The DDM provides a benchmark and a sanity check since we can compute its likelihood analytically. The Full-DDM provides us with a model for which analytical computations are still based on the analytical likelihood of the DDM, however evaluation is slowed by the necessity for numerical integration. This forms a first test for the speed of evaluation of our methods. For the Ornstein Uhlenbeck, Levy, Race and DDM with parameterized boundary models we cannot base our calculations on an analytical likelihood, but we can nevertheless perform parameter recovery and compare to other methods that utilize empirical likelihoods. The Ornstein Uhlenbeck Model adds state-dependent behavior to the diffusion while the Levy Model adds variation in the noise process and the Race Models expand the output dimensions according to the number of choices.

#### Full Drift Diffusion Model

The Full Drift Diffusion Model (Full-DDM) maintains the same specification for the driving SDE, but also allows for trial-to-trial variability in three parameters (***Ratcliff and McKoon, 2008***). We allow the drift rate *v* to vary trial by trial, according to a normal distribution, 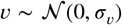, the non-decision-time *r* to vary according to a uniform distribution *τ* ~ **U**[−*ϵ*_*τ*_, *ϵ*_*τ*_] and the starting point *w* to vary according to a uniform distribution as well *w* ~ **U**[−*ϵ*_*w*_, *ϵ*_*w*_]. The parameter vector for the Full-DDM model is then *θ* = (*v, a, w, τ*, *σ*_*v*_, *ϵ*_*τ*_, *ϵ*_*w*_).

To calculate the FPTD for this model, we can use the analytical likelihood expression from the DDM. However, we need to use numerical integration to take into account the random parameters (***Wiecki et al., 2013***). This inflates execution time by a factor equivalent to the number of executions needed to compute the numerical integral.

#### Ornstein Uhlenbeck Model

The Ornstein Uhlenbeck model introduces a state-dependency on the drift rate *v*. Here *a*(*t, x*) = *v* + *g* * *x*, where *g* is an inhibition / excitation parameter. If *g* < 0 it acts as a leak (the particle is mean reverting). If *g* > 0 the particle accelerates away from the 0 state, as in an attractor model. At *g* = 0 we recover the simple DDM process. This leaves us with a parameter vector *θ* = (*v, a, w, τ, g*).

The corresponding SDE is defined as,

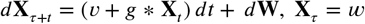

This model does not have an analytical likelihood function that can be employed for cheap inference (***Mullowney and Iyengar, 2006***). We discuss alternatives, other than our proposed methods, to simple analytical likelihoods later. For our purposes approximate inference is necessary for this model. ^3^.

#### Levy Flights

The Levy Flight (***Wieschen et al., 2020***; ***Reynolds and Rhodes, 2009***) model dispenses with the Gaussian noise assumption in that the incremental noise process instead follows an alpha-stable distribution 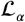. Specifically, we consider distributions 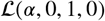 which are centered at 0, symmetric and have unitary scale parameter. These distributions have a first moment for *α* ∈ (1, 2], but infinite variance for *α* < 2. An important special case is *α* = 2, where 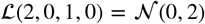. The parameter vector for this process is *θ* = (*v, a, w, τ, α*). We fix *a*(*t, x*) = *v* and *b*(*t, x*) = 1. The SDE is defined as,

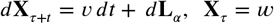

The Levy Flight is a flexible model used across disciplines for some of its theoretical optimality properties (***Wosniack et al., 2017***) despite not possessing closed-form FPTD’s. We add it here, as it is different from the other models under consideration; in principle, it could also capture decision-making scenarios in which there are sudden jumps in the accumulation of evidence (e.g., due to internal changes in attention). Its behavior is shaped by altering the properties of the incremental noise process directly.

#### Parameterized Collapsing Decision Bounds

We will consider variations of the DDM in which the decision boundary is not fixed but is time-varying (represented by a boundary parameter *a* with a parameterized boundary function *h*(*t*; *θ*_*h*_)). In such cases we augment the parameter vector *θ* with the set *θ*_*h*_ and drop *a*. Such variations are optimal in a variety of settings (for example, when there are response deadlines (***Frazier and Yu, 2008***) or distributions of trial-types with different difficulties (***Malhotra et al., 2018***; ***Palestro et al., 2018***)), and also better reflect the underlying dynamics of decision bounds within biologically inspired neural models (***O’Reilly and Frank, 2006***; ***Ratcliff and Frank, 2012***; ***Wiecki and Frank, 2013***). The boundary functions considered in the following are the Weibull bound (WEIBULL),

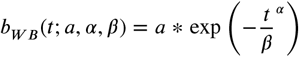

and the linear collapse bound (ANGLE),

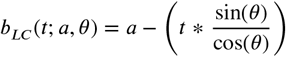

#### Race Models: N > 2

The Race Model departs from previous model formulations in that it has a particle for each of *N* choice options, instead of a single particle representing the evidence for one option over another. The function *f*_*E*_*i*__ (*t, 0*) now represents the probability of particle to be the first of all particle to cross the bound *a* at time *t*. We consider race models for which the drift and starting point can vary for each particle separately. Treating the boundary as a constant *a* leaves us with a parameter vector *θ* = (*v*_1_, …, *v*_*n*_, *a, w*_1_, …, *w_n_, ndt*). The SDE is defined, for each particle separately (or in vector form) as,

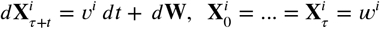

These models represent the most straightforward extension to a multi-choice scenario.

### MLP

#### Network Specifics

We apply the same simple architecture consistently across all example contexts in this paper. Our networks have 3 hidden layers, {*L*_1_, *L*_2_*, L*_3_} of sizes {100, 100, 120}, each using *tanh*(.) activation functions. The output layer consists of a single node with linear activation function.

#### Training Process

##### Training Hyperparameters

The network is trained via stochastic back-propagation using the Adam (***Kingma and Ba, 2015***) optimization algorithm. As a loss function, we utilize the huber loss (***Huber, 1992***) defined as,

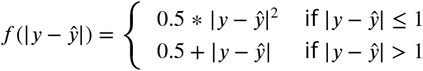

##### Training Data

We used the following approach to generate training data across all examples shown below.

First, we generate 100*K* simulations from the stochastic simulator (or model 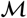)), for each of 1.5*M* parameter configurations. Since for the examples we consider, the stochasticity underlying the models are in the form of a stochastic differential equation (SDE), all simulations were conducted using the simple Euler-Maruyama method with timesteps *δt* of 0.001*s*. The maximum time we allowed the algorithms to run was 20*s*, much more than necessary for a normal application of the simulator models under consideration.

Based on these simulations, we then generate empirical likelihood functions using kernel density estimators (KDEs) (***Turner et al., 2015b***). KDEs use atomic datapoints {*x*_0_, …, *x*_*N*_} and reformulate them into a continuous probability distribution 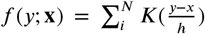, where we choose *K*(.) as a standard gaussian kernel 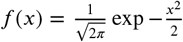, and *h* the so-called bandwidth parameter, is set by utilizing Silverman’s rule of thumb (***Silverman, 1986***). Where the data made Silverman’s rule inapplicable we set a lower bound on *h* as 10^−3^. Additionally we follow (***Charpentier and Flachaire, 2015***) in transforming our KDE to accommodate positive random variables with skewed distributions (in adherence to the properties of data resulting from the response time models forming our examples).

To ensure that the networks accurately learn likelihoods across a range of plausible data, for each parameter set we trained the networks by sampling 1000 datapoints from a mixture distribution with three components (mixture probabilities respectively {0.8, 0.1, 0.1}. The first component draws samples directly from the KDE-distributions. The second component is uniform on [0*s,* 20*s*], and the third component samples uniformly on [−1*s,* 0*s*]. The aim of this mixture is to allow the network to see, for each parameterization of the stochastic simulator, training examples of three kinds: (1) “Where it matters”, that is where the bulk of the probability mass is given the generative model. (2) Regions of low probability to inform the likelihood estimate in those regions (i.e. to prevent distortion of likelihood estimates for datapoints that are unlikely to be generated under the model). (3) Examples on the negative real line to ensure that it is learned to consistently drive likelihood predictions to 0 for datapoints close to 0.

The supervision signal for training has two components. For positive datapoints (reaction times in our examples), we evaluate the log likelihood according to our KDE. Likelihoods of negative datapoints were set to an arbitrary low value of 10^−29^ (a log likelihood of −66.79). 10^−29^ also served as the lower bounds on likelihood evaluations. While this constrains our accuracy on the very tails of distributions, extremely low evaluations unduly affect the training procedure. Since the generation of training data can easily be parallelized across machines, we simply front-loaded the data generation accordingly. We refer back to Figure 2 for a conceptual overview.

This procedure yields 1.5*B* labeled training examples on which we train the network. We applied early stopping upon a lack of loss improvement for more than 5 epochs of training. All model were implemented using Tensorflow (***Abadi et al., 2016***).

We note here that this amount of training examples is likely an overshoot by potentially one or more orders of magnitude. We did not systematically test for the minimum amount of training examples needed to train the networks. Minimal experiments we ran showed that roughly one tenth of the training examples lead to very much equivalent training results. Systematic minimization of the training data is left for future numerical experiments, since we don’t deem it essential for purposes of a proof of concept.

#### Sampling algorithms

Once trained, we can now run standard Markov Chain Monte Carlo schemes, where instead of an analytical likelihood, we evaluate *f*_*w*_(**x***, θ*) as a forward pass through the MLP. Figure 1 *B* schematically illustrates this approach (following the green arrows), and contrasts with currently applied methods (red arrows). We report multiple so-conducted parameter recovery experiments in the Results section and validate the approach first with models with known analytic likelihood functions.

Regarding sampling we utilized two MCMC algorithms, which showed generally very similar results. In contrast to the importance sampling algorithm used for the CNN (described below), MCMC methods are known for having trouble with multimodal posteriors. Running our experiments across algorithms was a safeguard against incorporating sampler specific deficiencies into our analysis. We however acknowledge that even more extensive experiments may be necessary for comprehensive guarantees. First, having an ultimate implementation of our method into the HDDM Python toolbox (***Wiecki et al., 2013***) in view, we use slice sampling (as used by the toolbox), specifically the step-out procedure following ***Neal (2003)***. Second, we used a custom implementation of the DE-MCMC algorithm (***Ter Braak, 2006***), known for being robust in higher dimensional parameter spaces. Our DE-MCMC implementation adds reflecting boundaries to counteract prob-lematic behavior when the sampler attempts to move beyond the parameter space, which is truncated by the (broad) range of parameters in which the MLP was trained. The number of chains we use is consistently determined as 5 * *θ*, five times the number of parameters of a given stochastic model. Samplers were initialized, by using slight perturbations of 5 maximum likelihood estimates, computed via differential evolution optimization (***Storn and Price, 1997***; ***Virtanen et al., 2020***). Since results were very similar across samplers, we restrict ourselves mostly to reporting results derived from the slice sampler, given that this sampler forms the back-end of the HDDM user interface we envision. ^4^.

#### Additional Notes

Note that we restricted parameter recovery for the MLP to datasets which distributed at least 5% of choices to the less frequently chosen option. This modest filtering accommodates the fact that such data-sets were also excluded form the training data for the MLP model, since they (1) present difficulties for the KDE estimator, (2) lead to generally less stable parameter estimates (i.e., it is not advisable to use diffusion models when choices are deterministic).

### CNN

#### Network Specifics

The CNN takes as an input a parameter vector *θ*, giving as output a discrete probability distribution over the relevant dataspace. In the context of our examples below the output space is of dimensions 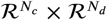, where *N*_*c*_ is the number of relevant choice alternatives, and *N*_*d*_ is the number of bins for the reaction time for each choice (*N*_*d*_ = 512 for all examples below). The network architecture consists of a sequence of three fully connected up-sampling layers, 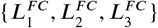, of respectively {64, 256, 1024} nodes. These are followed by a sequence of three convolutional layers 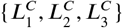 with 1 × 5 kernels, and a final fully connected layer with softmax activation. The network size was not minimized through architecture search, which along with other potential further speed improvements we leave for future research.

#### Training Process

For the CNN, we use 100*K* simulations from the stochastic simulator for each of 3*M* parameterizations, and bin the simulation outcomes as normalized counts into 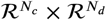 slots respectively (looking ahead to our examples, *N*_*c*_ concerns the number of choice outcomes, and *N*_*d*_ the number of bins into which the reaction time outcomes are split for a given simulator). The resultant relative frequency histograms (empirical likelihood functions) 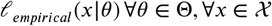, then serve as the target labels during training, with the corresponding parameters *θ* serving as feature vectors. For a given parameter vector *θ* the CNN gives out a histogram 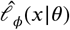, where *φ* are the network parameters. The network is then trained by minimizing the KL-divergence between observed and generated histograms,

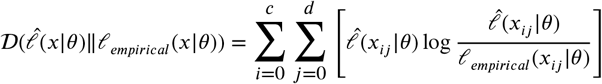

Training 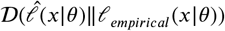, is not the only option. We note that it would have been a valid choice to train on 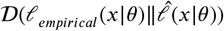 ***Minka, 2013***), or the symmetrized Kullback Leibler Divergence instead. Training results however were good enough for our present purposes to leave a precise performance comparison across those loss functions for future research, leaving room for further improvements.

As for the MLP, we use the Adam optimizer (***Kingma and Ba, 2015***), and implemented the network in Tensorflow (***Abadi et al., 2016***).

#### Sampling Algorithm

One benefit in using the CNN lies in the enhanced potential for parallel processing across large number of parameter configurations and datapoints. To fully exploit this capability, instead of running a (sequential) MCMC algorithm for our parameter recovery studies, we use iterated importance sampling, which can be done in parallel. Specifically, we use adaptive importance sampling based on mixtures of t-distributions, following a slightly adjusted version of the suggestions in (***Cappé et al., 2008***; ***Wraith et al., 2009***).

While importance sampling is well established, for clarity and the setting in which we apply it, we explain some of the details here. Importance sampling algorithms are driven by the basic equality,

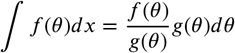

which holds for any pair of probability distributions such that *g*(*θ*) > 0*wheref* (*θ*) > 0. *f* (*θ*) is our posterior distribution, and *g*(*θ*) is the proposal distribution. We now sample *N* tuples *θ* according to *g*(*θ*), and assign each *θ*_*i*_ an importance weight 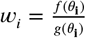.

To get samples from the posterior distribution, we sample with replacement the *θ* from the set {*θ*_0_, …, *θ*_*n*_}, with probabilities assigned as the normalized weights 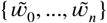. We note that importance sampling is exact for *N* → ∞. However for finite *N*, the performance is strongly dependent on the quality of the proposal distribution *g*(.). A bad match of *f* (.) and *g*(.) leads to high variance in the importance weights, which drives down performance of the algorithm, as commonly measured by the effective sample size (***Liu, 2008***),

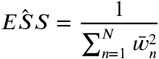

Iterated importance sampling uses consecutive importance sampling rounds, to improve the proposal distribution *g*(.). A final importance sampling round is used to get the importance sample we use as our posterior sample. Specifically, we start with a mixture of t-distributions *g*_0_(.), where 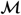 is the number of mixture components. Each component of *g*_0_(.) is centered at the MAP according to a optimization run (again we used differential evolution). The component-covariance-matrix is estimated by a numerical approximation of the Hessian at the respective MAP. Each round, based on the importance sample {**x**, **w**}_*i*_, we update the proposal distribution (to a new mixture of t-distributions), using the update equations derived in (***Cappé et al., 2008***).

As suggested by ***Cappé et al. (2008)***, convergence is assessed using the normalized perplexity statistic (the exponentiated Shannon entropy of the importance weights). For run this is computed as 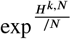, where 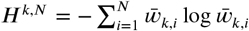.

To help convergence, we depart from the basic setup suggested in ***Cappé et al. (2008)*** in the following way. We apply an annealing factor *γ*_*k*_ = max 2^*z*−*k*,^ 1 *z* ∈ {1, 2, 4, …,}, so that for iteration *k* of the importance sampler, we are operating on the target 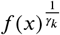. Smoothing the target during the first iterations helps with successfully adjusting the proposal distribution *g*(.). Figure 2 visualizes the CNN approach. Again, we emphasize that more numerical experiments using a larger variety of sampling algorithms are desirable, but are out of the scope for the current paper. ^5^

### Strengths and Weaknesses

In this section we clarify a few strengths and weaknesses of the two presented methods and their respective use cases. First, representing the likelihood function datapoint-wise as an MLP output, or globally via the CNN output histogram, affects the potential for parallelization. As exploited by the choice of sampler, the CNN is very amenable to parallelization across parameters, since inputs are parameter tuples only. Since the output is represented as a global likelihood histogram, the dataset-likelihood is computed as the summation of the elementwise multiplied of bin-loglikelihoods, with a correspondingly binned dataset (counts over bins). This has the highly desirable property of making evaluation cost (time) independent of dataset size. While the MLP in principle allows parallel processing of inputs, the data-point wise representation of input values ({*θ, x*}) makes the potential for cross-parameter parallelization dependent on data-set sizes.

While a single evaluation of the CNN is more costly, cross-parameter batch processing can make it preferable to the MLP. Second, the CNN has an advantage during training, where the representation of the output as a softmax layer, and corresponding training via minimization of the KL divergence, provides a more robust training signal to ensure probability distributions compared to the purely local one in which the MLP learns a scalar likelihood output as a simple regression problem. Third, and conversely, the MLP formulation exhibits a major advantage in its potential for trial-wise parameter estimates. Because it expresses likelihood point-wise, it allows one to estimate the impact of trial-wise regressors on model parameters during inference, without further training. It is for example common in the cognitive neuroscience literature to allow the cross-trial time-course of EEG, fMRI, or spike signals to be modelled as a trial-by-trial regressor on model parameters of e.g. drift diffusion models (***Wiecki et al., 2013***; ***Frank et al., 2015***; ***Cavanagh et al., 2011***; ***Herz et al., 2016***; ***Pedersen and Frank, 2020***). Another relevant example is the incorporation of latent learning dynamics. If a subject’s choice behavior are driven by reinforcement learning across stimuli, we can translate this into trial by trial effects on the parameterizations of a generative process model (***Pedersen and Frank, 2020***). These applications are implicitly enabled at no extra cost with the MLP method, while the trial by trial split multiplies the necessary computations for the CNN by the number *N* of data-points. We stress however that in general both the CNN, as well as the MLP can directly be used for hierarchical inference scenarios.

## A Parameter Recovery

Here we provide additional figures concerning parameter recovery studies. Table 1 summarizes the parameter-wise *R*^2^ between ground truth and the posterior mean estimates for each tested model and for each the CNN and MLP (where applicable) methods in turn. For the MLP, results are based on a reference run which used training data constructed from KDE empirical likelihoods utilizing 100*k* simulations each, and a slice sampler stopped with help of the Geweke Diagnostic. Results in the paper are based on slice samplers as well as Slice samplers, which explains why not all *R*^2^ values match exactly the ones found in other Figures. Our findings were however generally robust across samplers.

## B Manifolds / Likelihoods

We show the some examples of the likelihood manifolds for the various models that we tested.

DDM-SDV

Figure 13

ANGLE

Figure 14

WEIBULL

Figure 15

LEVY

Figure 16

ORNSTEIN

Figure 17.

FULL-DDM

Figure 18.

**Appendix 0 Table 1.**
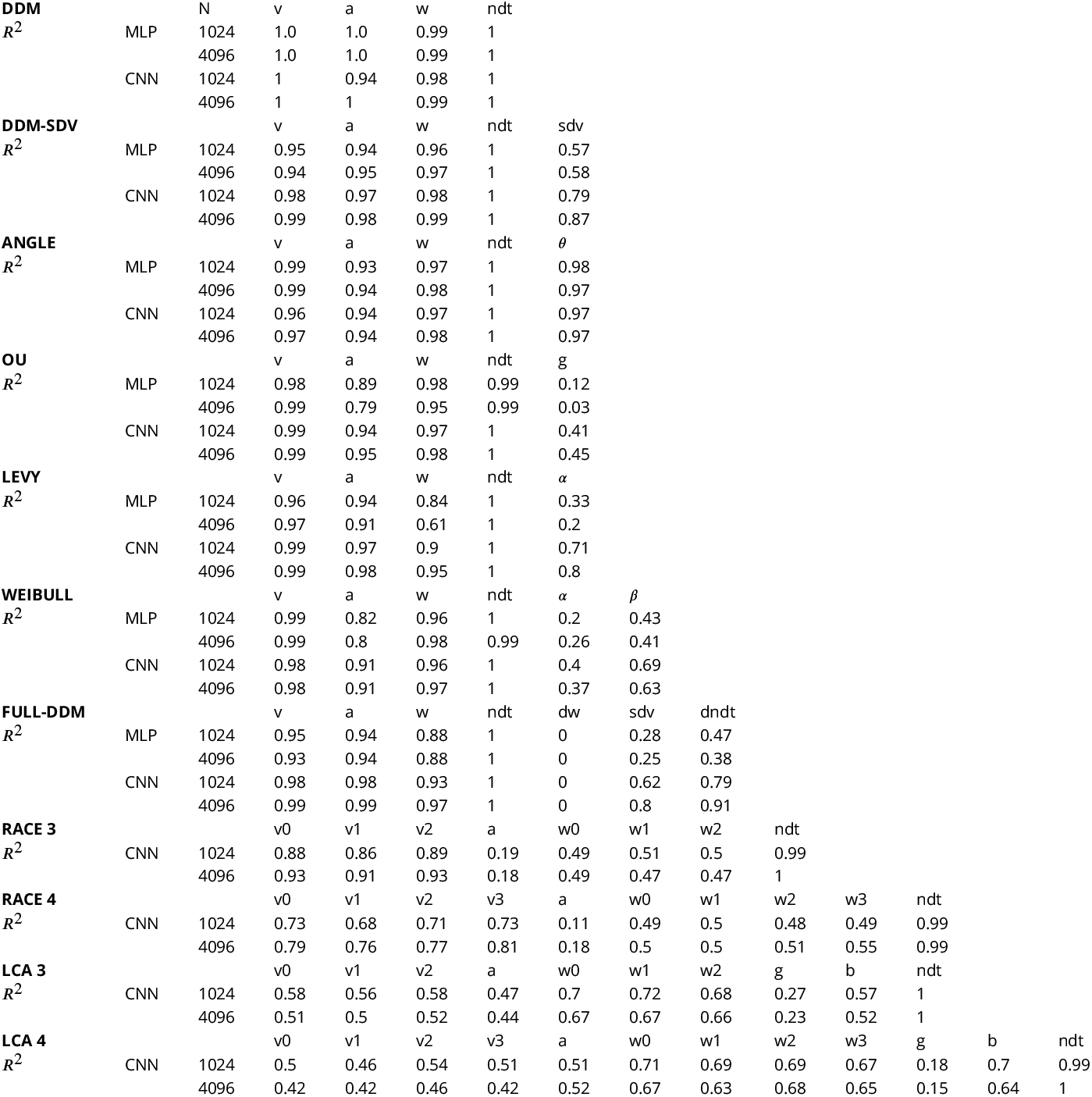
Parameter Recovery for a variety of test-bed models

**Appendix 0 Figure 13.**
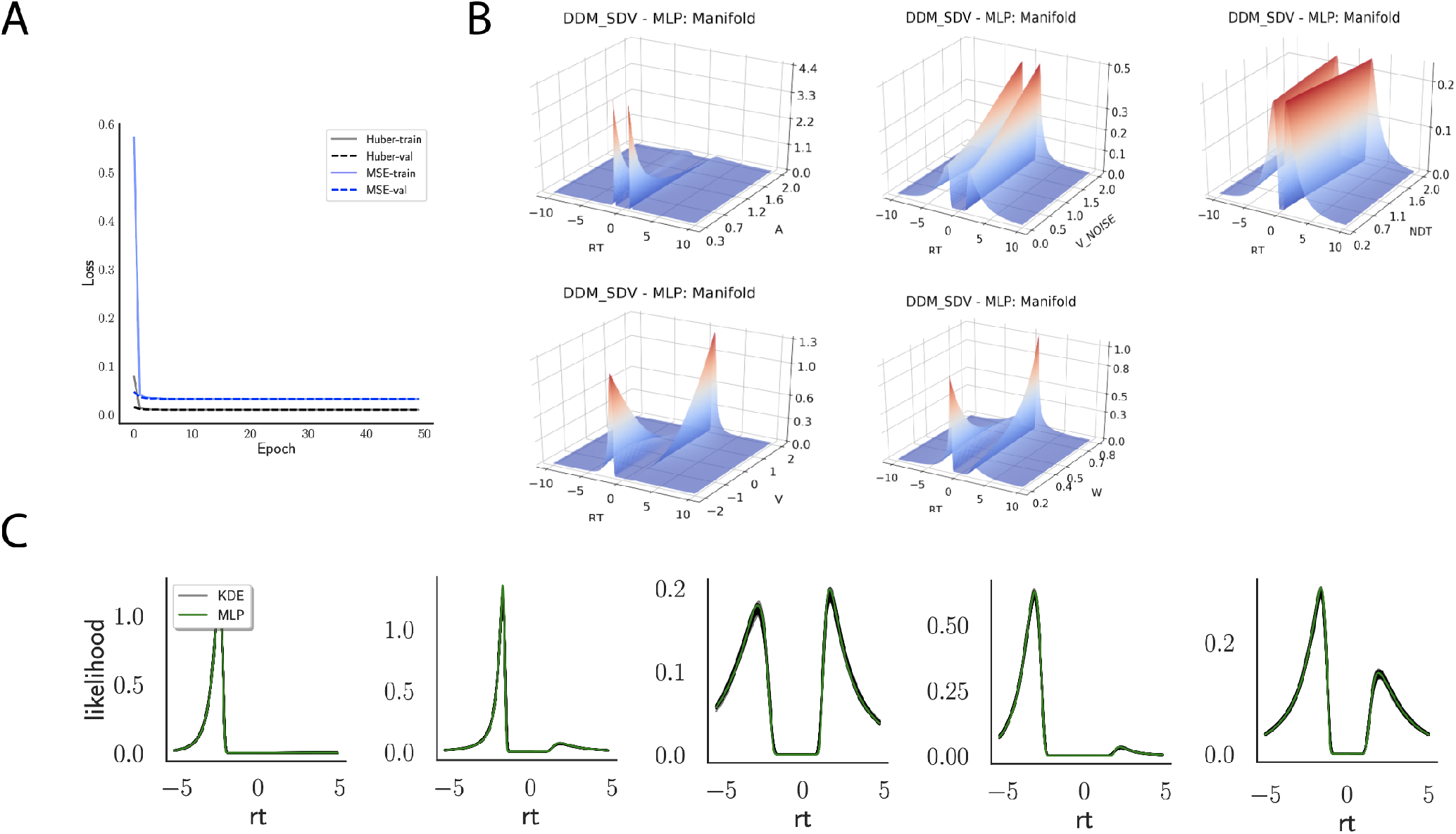
**A** Shows the training and validation loss for Huber as well as MSE for the DDM-SDV model. Training was driven by the Huber loss. **B** Illustrates the likelihood manifolds, by varying one parameter in the trained region. **C** shows MLP likelihoods in green, on top of a sample of 50 KDE-based empirical likelihoods derived from 20k samples each.

**Appendix 0 Figure 14.**
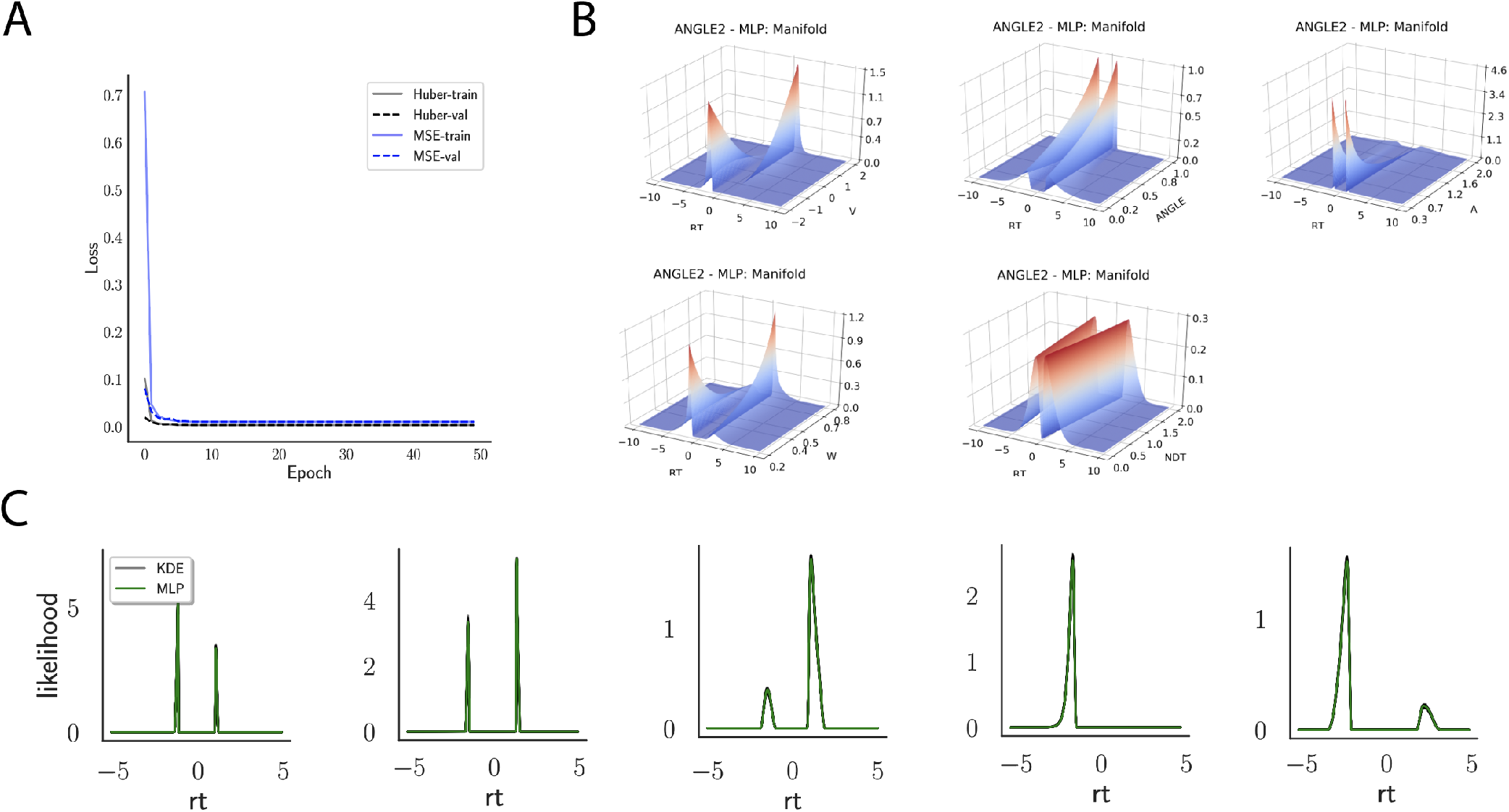
**A** Shows the training and validation loss for Huber as well as MSE for the ANGLE model. Training was driven by the Huber loss. **B** Illustrates the likelihood manifolds, by varying one parameter in the trained region. **C** shows MLP likelihoods in green, on top of a sample of 50 KDE-based empirical likelihoods derived from 20k samples each.

**Appendix 0 Figure 15.**
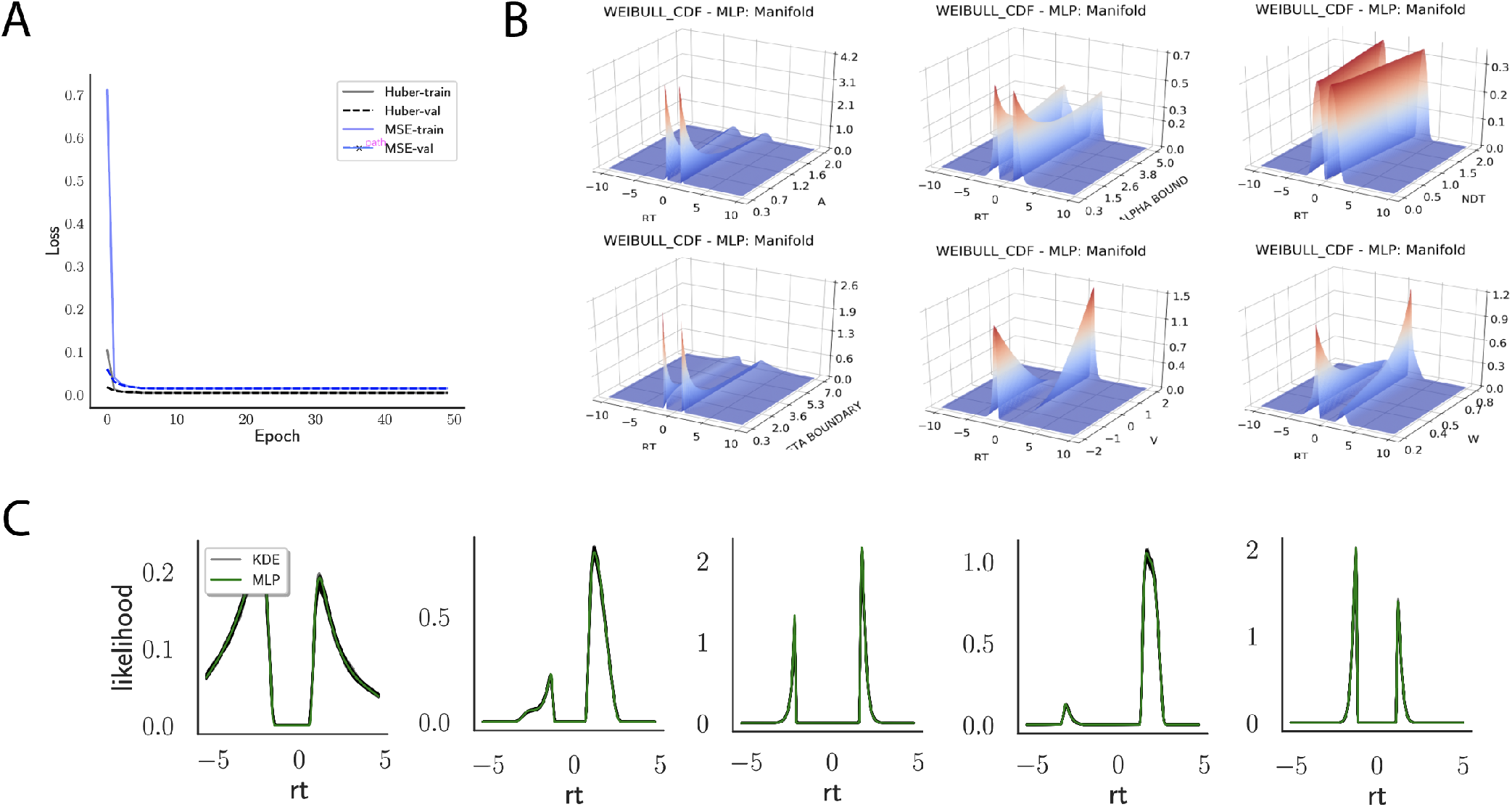
**A** Shows the training and validation loss for Huber as well as MSE for the WEIBULL model. Training was driven by the Huber loss. **B** Illustrates the likelihood manifolds, by varying one parameter in the trained region. **C** shows MLP likelihoods in green, on top of a sample of 50 KDE-based empirical likelihoods derived from 100k samples each.

**Appendix 0 Figure 16.**
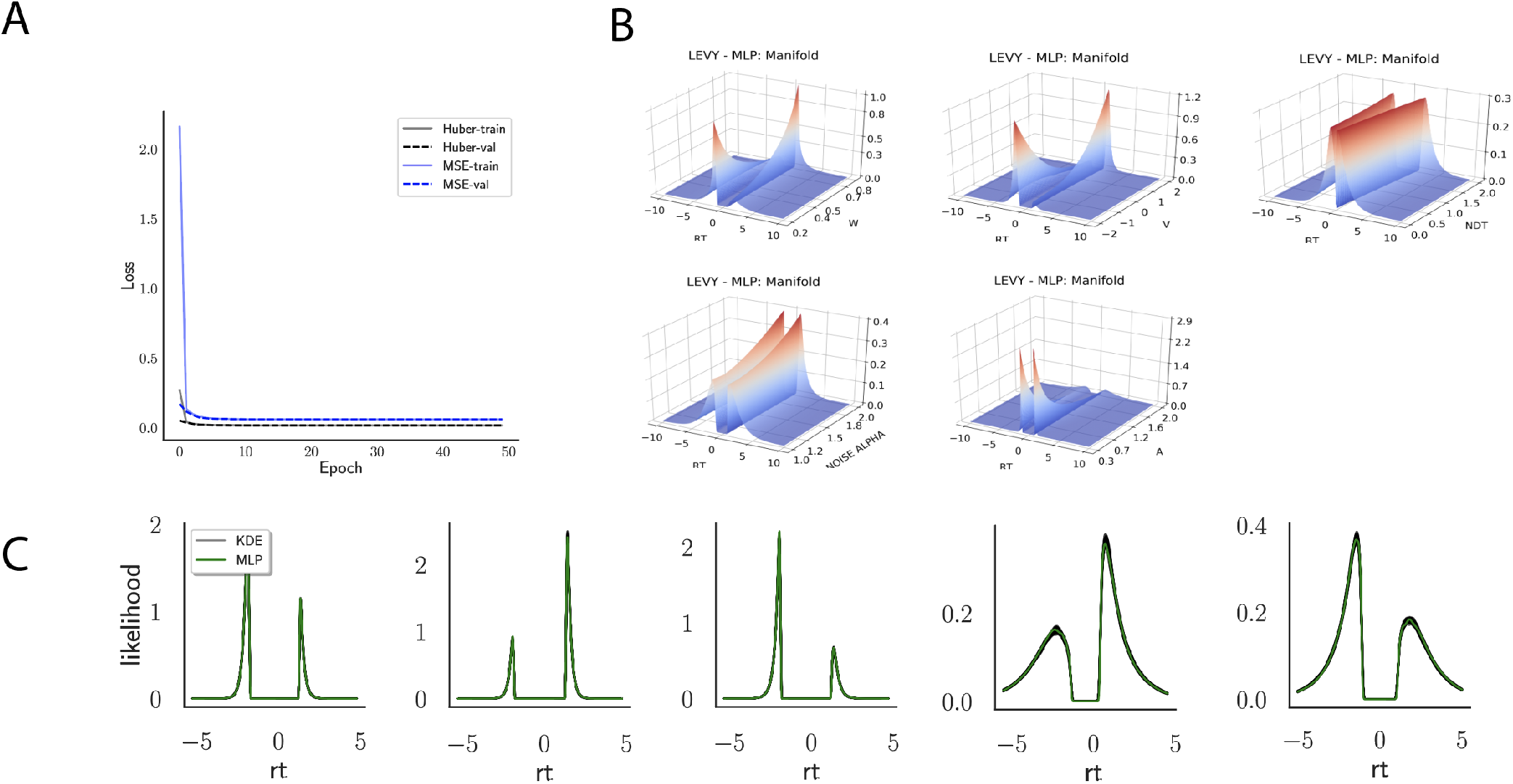
**A** Shows the training and validation loss for Huber as well as MSE for the Levy model. Training was driven by the Huber loss. **B** Illustrates the likelihood manifolds, by varying one parameter in the trained region. **C** shows MLP likelihoods in green, on top of a sample of 50 KDE-based empirical likelihoods derived from 100k samples each.

**Appendix 0 Figure 17.**
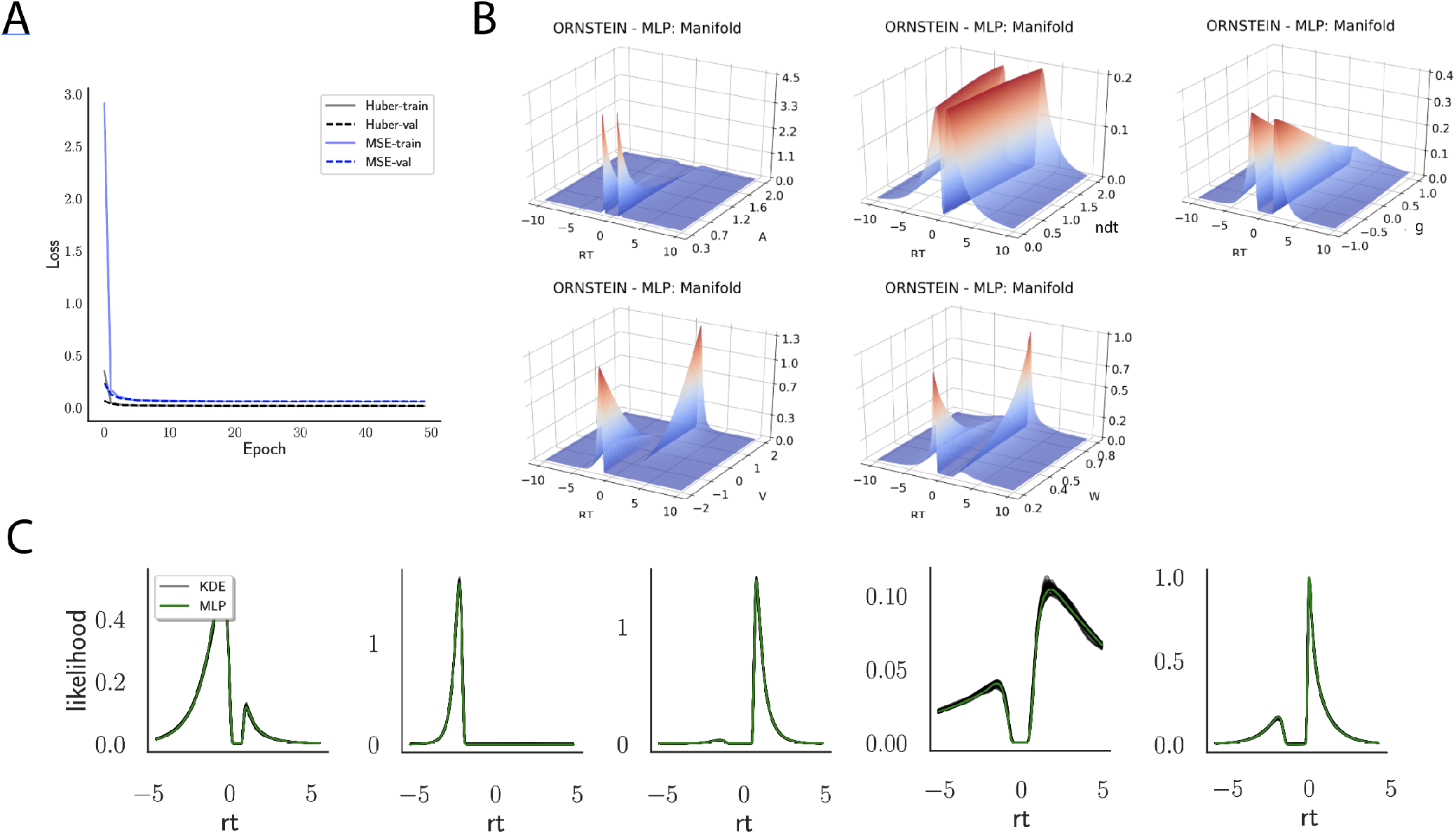
**A** Shows the training and validation loss for Huber as well as MSE for the ORNSTEIN model. Training was driven by the Huber loss. **B** Illustrates the likelihood manifolds, by varying one parameter in the trained region. **C** shows MLP likelihoods in green, on top of a sample of 50 KDE-based empirical likelihoods derived from 100k samples each.

**Appendix 0 Figure 18.**
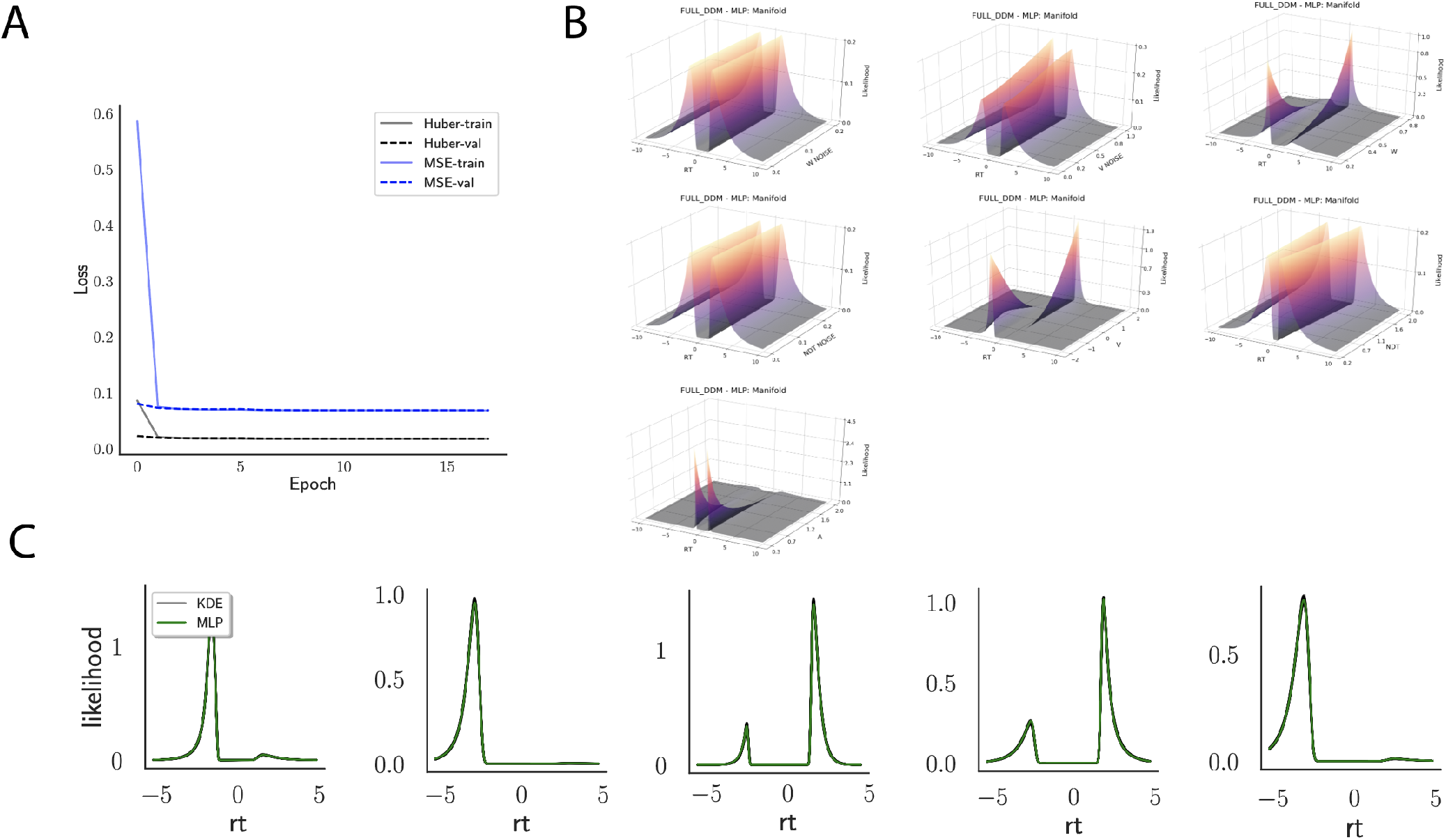
**A** Shows the training and validation loss for Huber as well as MSE for the Full-DDM model. Training was driven by the Huber loss. **B** Illustrates the likelihood manifolds, by varying one parameter in the trained region. **C** shows MLP likelihoods in green, on top of a sample of 50 KDE-based empirical likelihoods derived from 100k samples each

## Footnotes

1 We note two observations regarding sufficient statistics. First, a sufficient statistic is, in principle, simply a function of the data *f* ({*x*_1_, …, *x*_*n*_}) ⟼ {*s*_1_, …, *s*_*m*_}, where hopefully *m* << *n*. The identity map is one such function, whence we can consider the data itself as a special case (*s* can be *x*). Second, it is not common to have well-defined, low dimensional sufficient statistics for any given 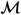.

2 Preliminary access to this interface and corresponding instrucitons can be found under https://github.com/lnccbrown/al-nets/tree/master/hddmnn-tutorial.

3 The Ornstein Uhlenbeck model is usually defined only for *g* < 0, our parameterization is strictly speaking a relaxation

4 Implementations of the MLP method, the samplers we used as well as the training pipeline can be found at https://github.com/lnccbrown/al-nets/tree/master/al-mlp.

5 Implementations of the CNN method, the samplers we used as well as the training pipeline can be found at https://github.com/lnccbrown/al-nets/tree/master/al-cnn.

